# Chromatin Acetylation Overrides Substrate Stiffness-Dependent Mechanoresponse and Lineage Specification in Human Mesenchymal Stem Cells

**DOI:** 10.1101/2022.01.04.474891

**Authors:** Rohit Joshi, Sourav Mukehrjee, Pooja Dharmambal Muralidharan, Tejas Suryawanshi, Pushpendra Yadav, Vedanshi Dharnidharka, Shobha Shukla, Abhijit Majumder

**Affiliations:** Department of Chemical Engineering, Indian Institute of Technology Bombay, Mumbai, India, 400076; Center for Research in Nanotechnology and Science, Indian Institute of Technology Bombay, Mumbai, India, 400076; Department of Metallurgical Engineering and Materials Science, Indian Institute of Technology Bombay, Mumbai, India, 400076

**Keywords:** Mesenchymal Stem Cells, substrate stiffness, mechanobiology, HDACi, HATi, chromatin remodelling, histone modification

## Abstract

Human mesenchymal stem cells (hMSCs) exhibit a stiffness dependent cellular phenotype and lineage bias. Cells on soft substrate display reduced cell spreading, less cellular traction, smaller and fewer focal adhesions and decreased actin stress fibre organization, whereas cells on stiff substrate exhibit the converse behaviour. Further, hMSCs preferentially undergo adipogenic differentiation on soft substrate and osteogenic differentiation on stiff substrate. These stiffness dependent changes are accompanied by global chromatin condensation/decondensation on soft/stiff substrates. Despite these well documented correlations, it remains unclear whether modulating chromatin condensation alone can be sufficient to reprogram stiffness dependent cellular phenotype. To answer whether chromatin organization acts as an active regulatory element in rigidity sensing, we induced chromatin condensation/decondensation in hMSCs cultured on soft/stiff polyacrylamide gels using histone deacetylase inhibitor (HDACi) and histone acetyltransferase inhibitor (HATi) respectively. We found that across conditions, phenotypic outcomes including nuclear localization of mechanosignalling protein YAP correlated strongly with chromatin compaction state rather than with substrate stiffness when the cells were presented with combination of signals. Mechanistically pharmacological inhibition of ERK phosphorylation attenuated chromatin dependent lineage specification while leaving chromatin compaction unchanged, indicating that ERK phosphorylation operates downstream of chromatin state. Collectively, these findings suggests an inside-out signalling mechanism in which chromatin compaction state reprograms the stiffness dependent mechanoresponse and fate decisions in hMSCs. Altogether, this works provides a fundamental framework for integrating mechanical and epigenetic cues in designing biomaterial microenvironment for controlling hMSCs fate in tissue engineering and regenerative medicine.

**Significance:** Rigidity sensing of the adherent mammalian cells are explained as outside-in signalling in which the proteins at the cell membrane boundary are the first to sense the rigidity of their microenvironment. The information then get transduced inside the cell finally reaching the nucleus and changing epigenetic landscape. This work demonstrates that altering the chromatin architecture by histone modification significantly changes the mechano-response of the cells, including the assembly of adhesion proteins. This work indicates a strong inside-out signalling cascade and suggest that chromatin state regulation can decouple lineage specification from substrate mechanics, providing a potential strategy to achieve desired stem cell lineage when matrix rigidity is not optimal for desired outcome.

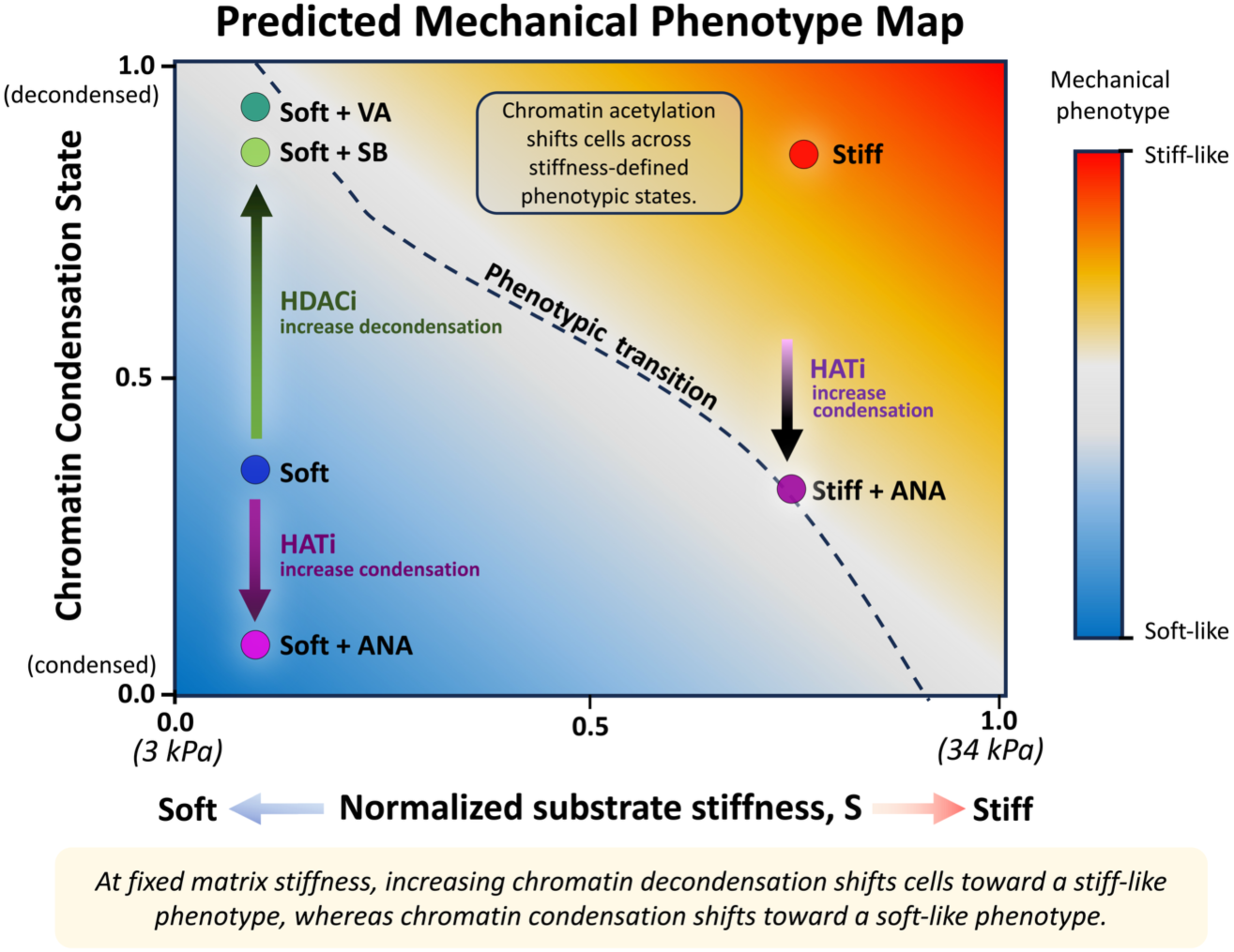

## Introduction

Mechanical cues from the extracellular matrix (ECM) play a central role in regulating cell behavior and tissue homeostasis. Among these mechanical cues, matrix stiffness is a major determinant of cell morphology, cytoskeletal organization, nuclear architecture, and lineage specification. Like many other adherent cells, human mesenchymal stem cells (hMSCs) when cultured on stiff substrate, they apply higher traction force which results into higher cell spreading, formation of stress fibers and larger focal adhesion as compared to their counterparts cultured on relatively softer materials [1, 2] This change in morphology due to stiffer environment gets transmitted to the nucleus through the acto-myosin cytoskeleton, the linker of nucleoskeleton and cytoskeleton complex (LINC), and nuclear lamina. This mechanical coupling alters nuclear morphology and is accompanied by changes in chromatin structure and gene regulation. Substrate rigidity is known to regulate hMSC lineage commitment, with stiffer substrates favoring osteogenic differentiation, whereas softer substrates promote adipogenic differentiation [3], [4], [5]. Previous studies have shown that hMSCs cultured on soft matrices exhibit relatively condensed chromatin and lower histone acetylation, whereas cells cultured on stiff matrices maintain a more decondensed and acetylated chromatin state [6] [7]. Such stiffness-dependent chromatin remodelling may provide a mechanism through which transient mechanical inputs are converted into persistent transcriptional and lineage-specific programs [8] [9]. Within this conventional outside-in framework, chromatin is often considered a downstream target of extracellular mechanics. However, accumulating evidence from literature indicates that chromatin is itself a mechanoresponsive structure [6] [10], [11], [12], [13], [14]. Nevertheless, it remains unclear whether chromatin organization is merely a consequence of matrix stiffness or it actively determines how cells interpret and respond to mechanical cues.

This distinction raises an important mechanobiological question: can experimentally altering chromatin compaction reprogram stiffness-dependent cellular phenotype without changing the mechanical properties of the substrate? More specifically, would inducing a decondensed, stiff-associated chromatin state in hMSCs cultured on soft substrates be sufficient to promote stiff-like mechanophenotypes and osteogenic bias? Conversely, would inducing chromatin condensation in cells cultured on stiff substrates promote soft-like cellular behaviour and adipogenic differentiation?

To address these questions, we modulated histone acetylation and chromatin compaction in hMSCs with valproic acid (VA), a histone deacetylase inhibitor (HDACi) [15], applied to hMSCs cultured on soft (3 kPa) substrates to increase global histone acetylation and promote chromatin decondensation. Conversely, anacardic acid (ANA), a histone acetyltransferase inhibitor (HATi) [16] was applied to cells cultured on stiff (34 kPa) substrates to reduce histone acetylation and promote chromatin condensation. Decondensing chromatin was observed to induce stiff-like cellular phenotypes in hMSCs cultured on soft substrates, characterized by increased spreading, focal-adhesion maturation, actin organization, traction-force generation, nuclear YAP localization and osteogenic differentiation. Conversely, chromatin condensation promoted several soft-like characteristics in cells cultured on stiff substrates.

Together, these findings identify histone acetylation-dependent chromatin compaction as an active regulatory layer in hMSC mechanobiology. Rather than functioning solely as a downstream consequence of extracellular mechanics, chromatin state can reprogram how hMSCs interpret matrix stiffness and coordinate mechanotransduction with lineage specification.

## Material and methods

### Substrate preparation

Polyacrylamide gels having elastic modulus of ∼3 kPa and ∼34kPa were prepared by crosslinking 40% poly-acrylamide and 2% bis-acrylamide solution. Protocol for substrate preparation and Elastic modulus value was adopted form previously reported work [17]. Briefly, the gel solution for desired stiffness (∼3 kPa and 34 kPa, Table S1) was mixed with ammonium per sulphate (1:100) and TEMED (1:1000) and a drop of 130µL was placed between two glass coverslips, one coated with 3-APTMS (Sigma) and other with hydrophobic coating. After polymerization, hydrophobic coverslip was removed. The gel was coated with type I collagen (25 µg/mL) (Invitrogen; A1048301) using Sulfo-SANPAH (G Biosciences BC38) based conjugation and kept at 4°C overnight as previously described [18].

### Cell culture

Bone marrow-derived hMSCs were purchased from Lonza (Cat.No. #PT-2501). hMSCs were cultured in low glucose Dulbecco’s Modified Eagle Medium (DMEM) (HiMedia; AL006) supplemented with 16% FBS (HiMedia; RM9955), 1% Antibacterial-Antimycotic (HiMedia; A002) and 1% Glutamax (Gibco; 35050) under humidified conditions at 37°C with 5% CO_2_. The cells were trypsinized with TrypLE™(Gibco; 12604021) at 70% confluency. The cells were seeded on polyacrylamide gels with 2000 cells/cm^2^ seeding density in 50 µL of media and flooded after 45 min when the cells were attached.

### Treatment with HDACi and HATi

To check the effect of HDACi and HATi on soft substrate 0.5mM of valproic acid (VA, PHR 1061) and 30 µM of anacardic acid (ANA, Sigma A7236) were added respectively, to the growth media at the time of flooding. For differentiation experiments, VA and ANA were added with differentiation media. For the experiments with sodium butyrate (SB, B5887), 0.5mM of SB was added in the similar manner.

### Differentiation assays

hMSCs were seeded at 2000 cells/cm^2^ in a 12-well culture plate and maintained in growth medium for 24h after which the media was replaced by respective differentiation media. Adipogenic (Invitrogen, A10410) and osteogenic (Invitrogen, A10069) differentiation kits were used. Cells were incubated for 9 days in adipogenic and 14 days for osteogenic induction media before quantitative assays. Differentiation media change was replenished after every 48h. After the completion of incubation, cells were fixed with 4% paraformaldehyde (PFA) for 30 min at room temperature followed by staining with Oil Red O (ORO, Sigma O0625) for adipogenic differentiation. After incubating with staining solution for 20 min samples are washed thrice with DPBS. Images were captured for quantitative analysis using EVOS inverted microscope (Life Technologies) in bright-field channel. ORO quantification was normalized by the total cells per view in percentage calculation.

### Immunofluorescence staining

Cells were fixed with 4% PFA in Dulbecco’s phosphate buffer saline (DPBS) for 15 min at room temperature (RT) then washed with DPBS thrice. Cells are then permeabilized with permeabilizing buffer (0.5% Triton X-100-Sigma Aldrich in DPBS) for 10 min and blocked with 4% Bovine serum albumin (BSA) in DPBS for 30 min to minimize non-specific protein binding. Anti-YAP (1:500, rabbit, Abcam, Cat. No. 52771), anti-PPAR-γ (1:500, rabbit, Abcam, Cat. No. 59256), anti-RUNX2 (1:500, rabbit, Abcam, Cat. No. 23981), anti-p-ERK (1:500, rabbit, Abcam, Cat. No. 65142), anti-OPN (1:500, rabbit, Abcam, Cat. No. 8448), anti-AcK (1:500, rabbit, ABCAM, Cat. No. 190479) and anti-pMLC (CST 3675S) primary antibodies in 4% BSA were added to the samples and incubated overnight at 4°C. Primary antibodies were removed and samples are rinsed with DPBS two times for 10 min. Samples were then incubated at room temperature with secondary antibodies (1:500, donkey anti-rabbit AlexaFlour 568 Cat. No. 75470, goat anti-rabbit AlexaFlour 488 Cat. No. 411034) phalloidin (1:400, AlexaFlour 532, Cat. No. A22282 and AlexaFlour 488, Cat. No. A12379) with Hoechst 33342 (Cat. No. H3570) of dilution 1:5000 in 4% BSA for 2 h at room temperature. Thereafter the samples were rinsed twice with DPBS followed by mounting of samples on glass slide with mounting media (ProLong, Thermo Fisher) and sealed with Eu-kit (Sigma–Aldrich). All samples were imaged at 63X (oil) magnification using laser scanning Confocal Microscope (LSM, Carl Zeiss). Z stack were acquired and maximum intensity projection from 3D stacks were used for fluorescence intensity calculation.

For vinculin staining hMSCs are fixed with ice cooled mixture of 1:1 (v/v) (4% PFA: permeabilizing buffer (1% Triton-X-100-Sigma Aldrich) for one minute on ice. Samples are rinsed with cytoskeleton stabilizing buffer (CSB) (60 mM PIPES, 27 mM HEPES, 10 mM EGTA, 4 mM magnesium sulphate, pH 7) and fixed again with 4% PFA in ice for 5 min. After fixing, samples were rinsed with CSB and blocked with 4% BSA supplemented with 0.5% Triton-X-100 for 30 min on ice. Anti-vinculin (1:500, rabbit Abcam Cat. No. ab129002) primary antibody in 4% BSA were added to the samples and incubated overnight at 4°C. Primary antibody was removed and samples are rinsed with CSB two times for 10 min. Samples were then incubated at room temperature with secondary antibody (1:500, goat anti-rabbit AlexaFlour 488 Cat. No. 411034), phalloidin (1:400, AlexaFlour 532, Cat. No. A22282) with Hoechst 33342 (Cat. No. H3570) of dilution 1:5000 in 4% BSA for 2 h at room temperature. All immunostained samples were imaged as previously mentioned.

### Analysis of chromatin condensation parameter (CCP)

DAPI stained nuclei were imaged with laser scanning confocal microscope (Carl Zeiss LSM 780) at 63X (oil). For analysis, individual nuclei were cropped using FIJI (Image J) to get only one nucleus per image. CCP was calculated using a MATLAB script as previously described [19] in which a gradient-based Sobel edge detection algorithm was used to measure the edge density of individual nuclei.

### Traction force microscopy (TFM) and Actin anisotropy

Gels of 3 kPa were made on 22 × 22 mm^2^ coverslips, once gels were solidified, 25µL drop of 3 kPa solution having 1 µm fluorescent beads (Fluka with a final concentration of 1:50) was added on the hydrophobic plate and then the solidified gel was inverted into it and allowed to solidify. Gels were then treated with Sulfo-SANPHA and coated with collagen I as mentioned above. Approximately 1000 cells were seeded per coverslip. After 24 h of cell seeding, cells were lysed using 100µL of 1% Triton-X in 2mL of complete media. Images of the stressed (before lysing) and unstressed (after lysing) gels were captured by the EVOS FL Auto cell imaging system (Life Technologies). An average of 10 cells was analysed per gel. The code from J.P.Butler was used to calculate the traction force [20]. Actin anisotropy which measures how well the filaments are ordered was measured with the ImageJ plugin FibrilTool which uses the concept based on nematic tensor [21]. The randomly aligned fibres give value close to 0 and highly aligned fibres give value close to 1.

### Statistical analysis

For statistical analysis were performed on triplicate samples (cell or nuclei) n>30 using random sampling from each sample. The images were analysed in FIJI (ImageJ) and the data was plotted using OriginLab software (IIT Bombay License). The data is represented as mean ± standard deviation unless mentioned otherwise. Statistical significance was checked by performing Student’s t-test and p value of <0.05 were considered as significant. (p<0.05 is indicated as *, p<0.01 is indicated as **, p<0.001 is indicated as ***, and non-significant is indicated as ns).

## Results

### 1. HDACi and HATi influence chromatin condensation and histone acetylation state in hMSCs

Histone acetylation is a major regulator of chromatin compaction in hMSCs (Fig. 1A). To determine whether chromatin packaging can be regulated independent of substrate rigidity, the global acetylation state of histone was modulated using HDACi (VA) and HATi (ANA). The global histone acetylation was quantified by immunostaining the hMSCs, cultured on substrates of soft (3 kPa) and stiff (34 kPa) polyacrylamide gels in the absence/presence of VA and ANA (Fig. 1B-E, S2). The global histone acetylation status of the stiff-cultured cells was found to be ∼ 2 times higher compared to soft-cultured cells (Fig 1F). Furthermore, VA treatment increased histone acetylation in soft cultured cells to approaching levels measured in stiff cultured cells (Fig. 1F). Conversely, ANA treatment reduced histone acetylation in stiff-cultured cells closely matching the soft condition (Fig. 1F), indicating the histone acetylation status can be modulated independent of substrate stiffness.

**Fig. 1.**
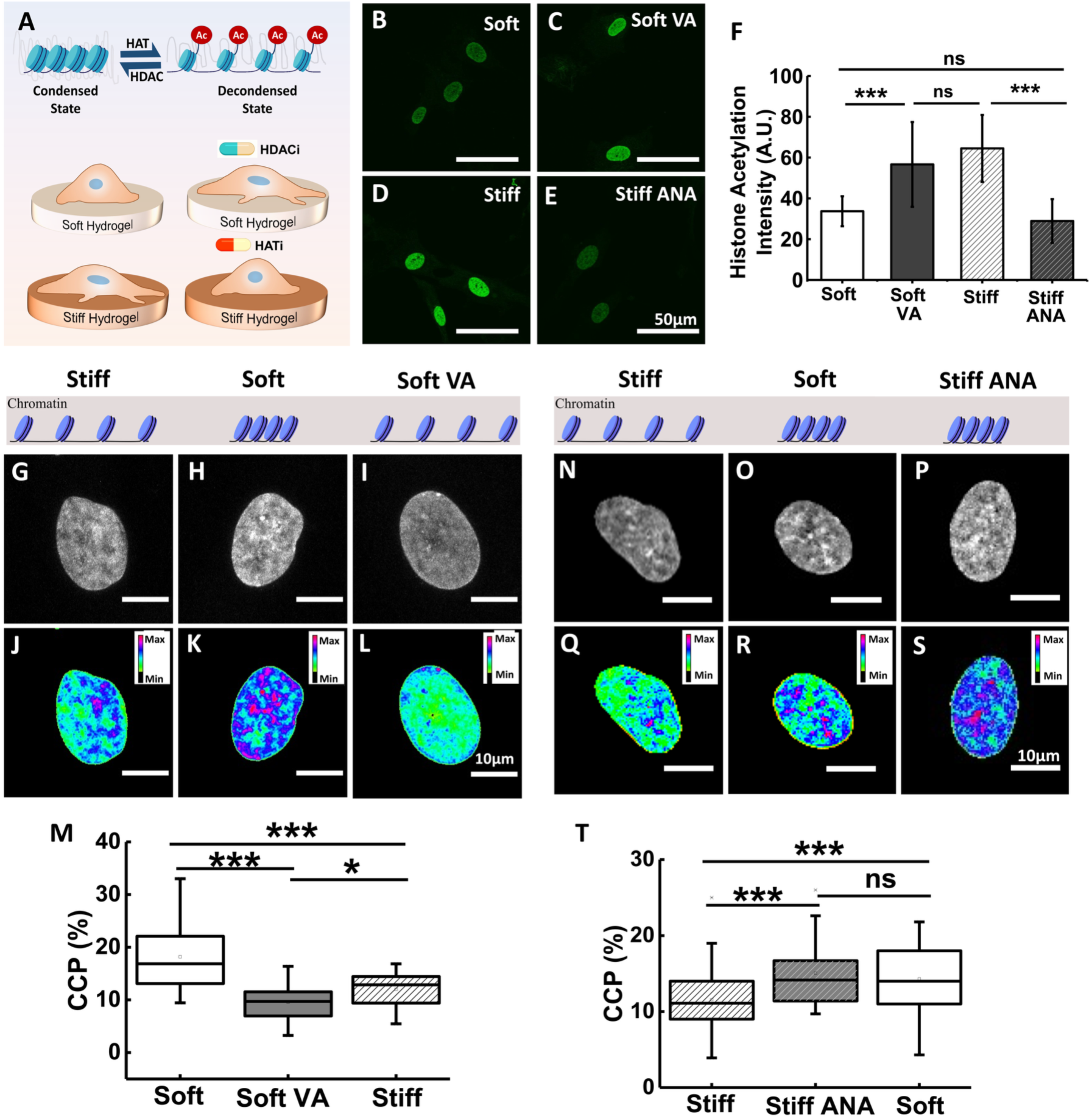
Influence of substrate stiffness and chromatin modifiers on chromatin condensation and histone acetylation. (A) Schematic showing histone acetylation mediated chromatin condensation/decondensation in response to HATi/HDACi treatment for the cells cultured on soft (3 kPa), soft + HDACi (VA), stiff (34 kPa) and stiff + HATi (ANA) respectively. (B-E) Representative images showing the average intensity of acetylated lysine residues in histones (green) for cells cultured on soft, soft+VA, stiff, and stiff+ANA respectively, scale bar 50 µm. (F) Quantification of the acetylated histone intensity has been shown. (G-I) Representative images of DAPI-stained nuclei of the hMSCs cultured on soft (3kPa), soft + VA and stiff (34kPa). (J-L) The corresponding heatmap of DAPI intensity. Warmer colours represent relatively more condensed chromatin, scale bar 10 µm. (M) Chromatin condensation parameter (CCP), defined as the percentage of highly DAPI-stained regions to the total nuclear area, for nuclei from soft, soft + VA, and stiff. (N-P) Schematic and DAPI-stained nuclei showing changes in chromatin condensation for the cells from stiff (34 kPa), stiff + ANA and soft (3 kPa). (Q-S) The corresponding heatmap of DAPI intensity. (T) Chromatin condensation parameter (CCP) for nuclei from stiff, soft, and stiff + ANA.

Further, to establish the baseline relationship between substrate stiffness and chromatin packing, chromatin compaction was quantified using chromatin condensation parameter (CCP). Briefly, high-magnification images of DAPI-stained nuclei in the hMSCs cultured on 3 kPa substrates with and without VA treatment were acquired (Fig. 1G–I). The bright DAPI-stained regions indicate densely packed chromatin (Fig. 1J-L). Consistent with the prior reports [13], we observed that hMSCs cultured on soft hydrogels had significantly higher chromatin condensation (∼ 2 times) (Fig. 1G-I, J-L, and M) compared to that on the stiff hydrogel.

Next, to remodel the chromatin packaging, hMSCs on the soft gel were treated with VA (0.5mM) for 24 h which increases global histone acetylation and promotes chromatin decondensation. The concentration of VA was chosen at 0.5 mM, as this condition demonstrated significant increase in histone acetylation and reduction in CCP compared to untreated cells on soft gels (Fig. S1), without affecting cell viability or proliferation. At this concentration, VA reduced chromatin condensation of cells on soft gels to the level comparable to that observed on stiff gels (Fig. 1J-M, S2) while the effect of VA persisted even 24 h after washout (Fig. S3). To confirm that the effect is not VA specific hMSCs were treated with another HDAC inhibitor, sodium butyrate (SB) and showed similar changes in global histone acetylation and CCP (Fig. S4), indicating the interpretation that chromatin decondensation on soft substrate is driven by HDAC inhibition rather than off target effect.

Similarly, to check whether chromatin compaction state on stiff substrate can be shifted towards a soft like condensed state, hMSCs cultured on stiff gels were treated with HATi (ANA), which reduced histone acetylation and promotes chromatin condensation (Fig. 1N-S). ANA treatment was found to increase the chromatin condensation ∼1.5 times on stiff-cultured cells (Fig. 1T).

While VA treatment reduced the CCP of hMSCs cultured on soft substrates similar to the level of stiff substrates (Fig. 1G-M), ANA treatment increased the CCP of hMSCs cultured on stiff substrates to levels similar to those observed on soft substrates (Fig. 1N-T). Together, these results indicate that pharmacological modulation of histone acetylation is sufficient to shift chromatin compaction to states associated with soft or stiff substrate, independent of substrate mechanics.

### 2. Chromatin remodelling overrides stiffness dependent cell spreading, focal adhesion and actin stress fibre formation

One of the earliest and most robust observable behaviour of cell is differential cell spreading, with cells exhibiting smaller spreading area on soft substrate and larger projected area on stiffer matrix [22], [23]. Hence, we investigated whether chromatin remodelling modulates mechanoresponsive phenotype of hMSCs cultured on soft and stiff substrate. Treatment of hMSCs with VA on 3 kPa gels resulted in significant increase in cell spreading (∼45%) compared to the untreated control as shown in (Fig. 2A-C). In addition to increased spreading, VA treated cells showed more membrane protrusions (Fig. S5), indicating enhanced cytoskeleton engagement with the substrate despite soft matrix.

**Fig. 2.**
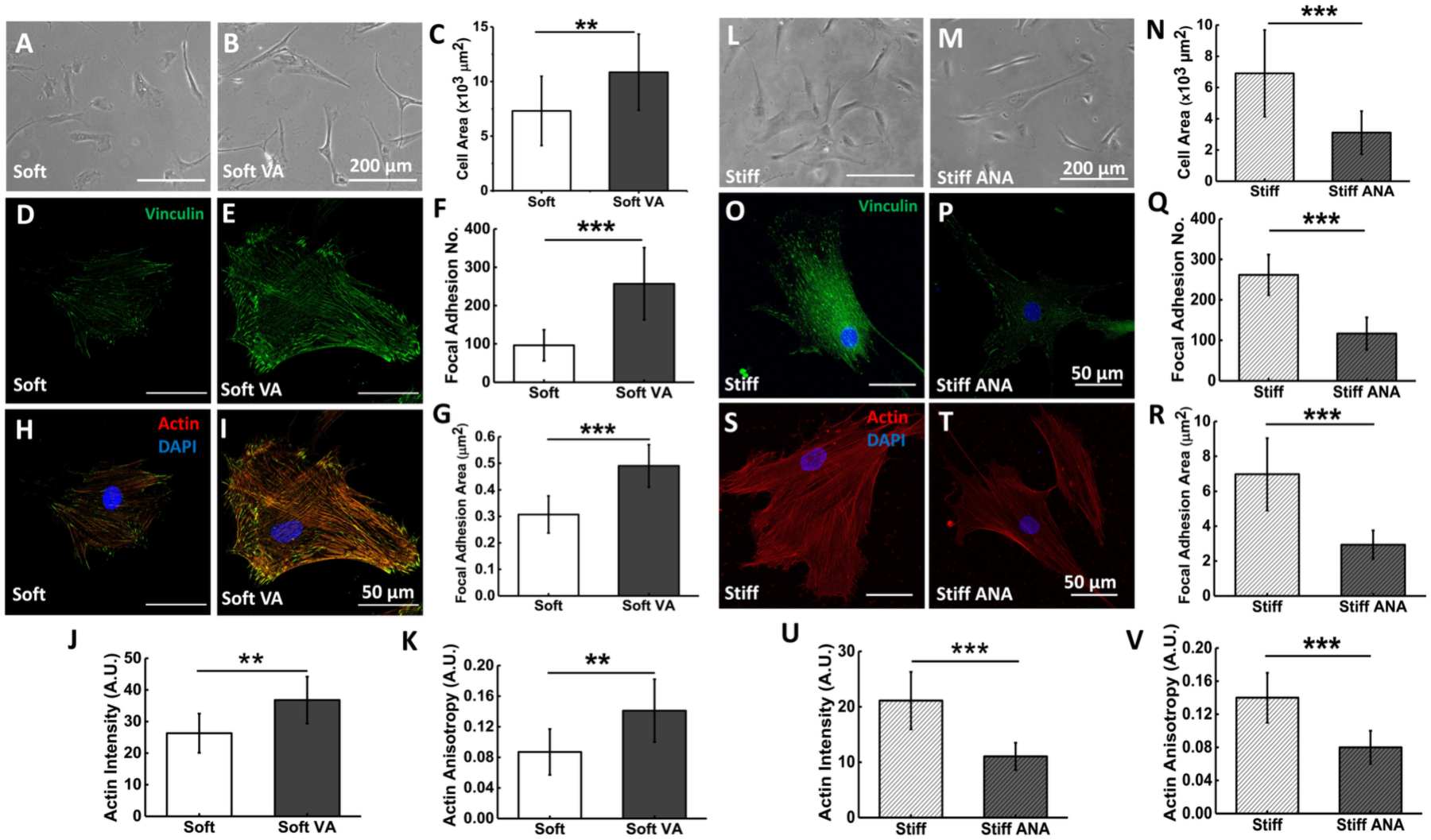
Chromatin remodelling overrides stiffness dependent cell spreading, focal adhesion and actin stress fiber formation. Representative phase contrast images of hMSCs cultured on soft (3 kPa) gel (A) without and (B) with VA (HDACi) has been shown. (C) The cell area of hMSCs has been quantified and shown, n=100, scale bar: 200 µm. (D and E) Representative images of hMSCs on 3 kPa gel showing immunostaining for vinculin (green) with and without VA respectively, scale bar: 50 μm. (F and G) The quantification of average focal adhesion number/cell and focal adhesion area has been shown for the cells cultured on soft substrate in presence and absence of VA. (H and I) hMSCs on 3 kPa gel were immunostained for actin (red) and DAPI (blue) with and without VA respectively. (J-K) The quantification of actin intensity and actin anisotropy for the cells cultured on soft substrate in presence and absence of VA, (L and M) Representative phase contrast images of hMSCs cultured on stiff (34 kPa) gel without and with ANA (HATi), scale bar: 200 μm. (N) The quantification of the cell area has been represented. (O and P) Representative images of hMSCs on 34 kPa gel showing immunostaining for vinculin (green) and DAPI (blue) with and without ANA respectively, scale bar: 50 μm. (Q and R) The quantification of average focal adhesion area and focal adhesion number/cell has been shown for the cells cultured on stiff substrate in presence and absence of ANA. (S and T) The representative images of hMSCs on 34 kPa gel, immunostained for actin (red) and DAPI (blue) with and without ANA respectively, scale bar: 50 μm. (U-V) The quantification of actin intensity and actin anisotropy for the cells cultured on stiff substrate (34 kPa) in presence and absence of ANA.

As cell spreading is known to be strongly coupled with formation of matured focal adhesions (FA) and actin fibre assembly [24], we examined the effect of VA on these two critical morphometric features by staining FA protein (vinculin) and F-actin, respectively. The cells treated with VA on soft substrates exhibited increment in average area per FA by ∼ 60% compared to untreated control cells indicating maturation of adhesion points (Fig. 2D-F). The VA treated cells also showed significantly increased number of FAs/cell as compared to the untreated control cells (from 94 ± 40 to 256 ± 94) (Fig. 2G). Parallelly, VA treated cells also showed a significant increase in F-actin intensity and formation of prominent stress fibres (Fig. 2H-I, J-K & S3), a phenotype typically associated with cells cultured on stiff substrate as compared to soft. To verify that the effect was not VA-specific, we used another histone deacetylase inhibitor, sodium butyrate (SB) [25], and found a similar outcome (Fig. S6). Furthermore, to rule out ECM specific contributions, we changed the ECM from collagen I to fibronectin and found the similar results for cell spread area, acetylation and chromatin condensation parameter in presence of VA (Fig. S7).

Similarly, to check whether inducing chromatin condensation on stiff substrate could restore soft like cellular phenotype, hMSCs cultured on stiff substrate were treated with ANA. Consistent with the observation of VA treatment on soft, it was observed that compacting chromatin on stiff substrate reduced cell spread area by ∼58% (Fig. 2L-N), average FA number/cell (from 261 ± 50 to 116 ± 40) and average FA area by approximately 2.3 folds (Fig. 2O-R), F-actin expression (from 22.2 ± 5.7 A.U. to 11.8 ± 2.4 AU), and actin anisotropy (from 0.14 ± 0.03 to 0.08 ± 0.02) (Fig. 2S-V) indicating a less contractile phenotype typical of cells cultured on soft substrates.

Together, these results demonstrate that chromatin compaction state exerts dominant control over cell spreading, focal adhesion maturation, and actin cytoskeleton organization, effectively overriding substrate dependent mechanoresponse.

### 3. Chromatin remodelling overrides rigidity dependent cellular tractions in hMSCs

Cell generated contractility or traction plays a critical role in mechanosensing. Majority of the cellular mechanoresponse is mediated via this actomyosin driven force [26], [27]. Hence, to answer this question, cellular traction for hMSCs cultured was estimated on soft PAA hydrogel (*E =* 3 kPa) in presence and absence of VA using traction force microscopy (TFM) (Fig. 3A). It was observed that treatment with VA significantly increased the traction force generated by hMSCs cultured on soft substrates (from ∼ 152 ± 32 Pa to ∼ 235 ± 39 Pa) [28] (Fig. 3B-D). Additionally, treatment with SB produced similar effects, confirming that the observed response was not specific to VA (Fig. S8).

**Fig. 3.**
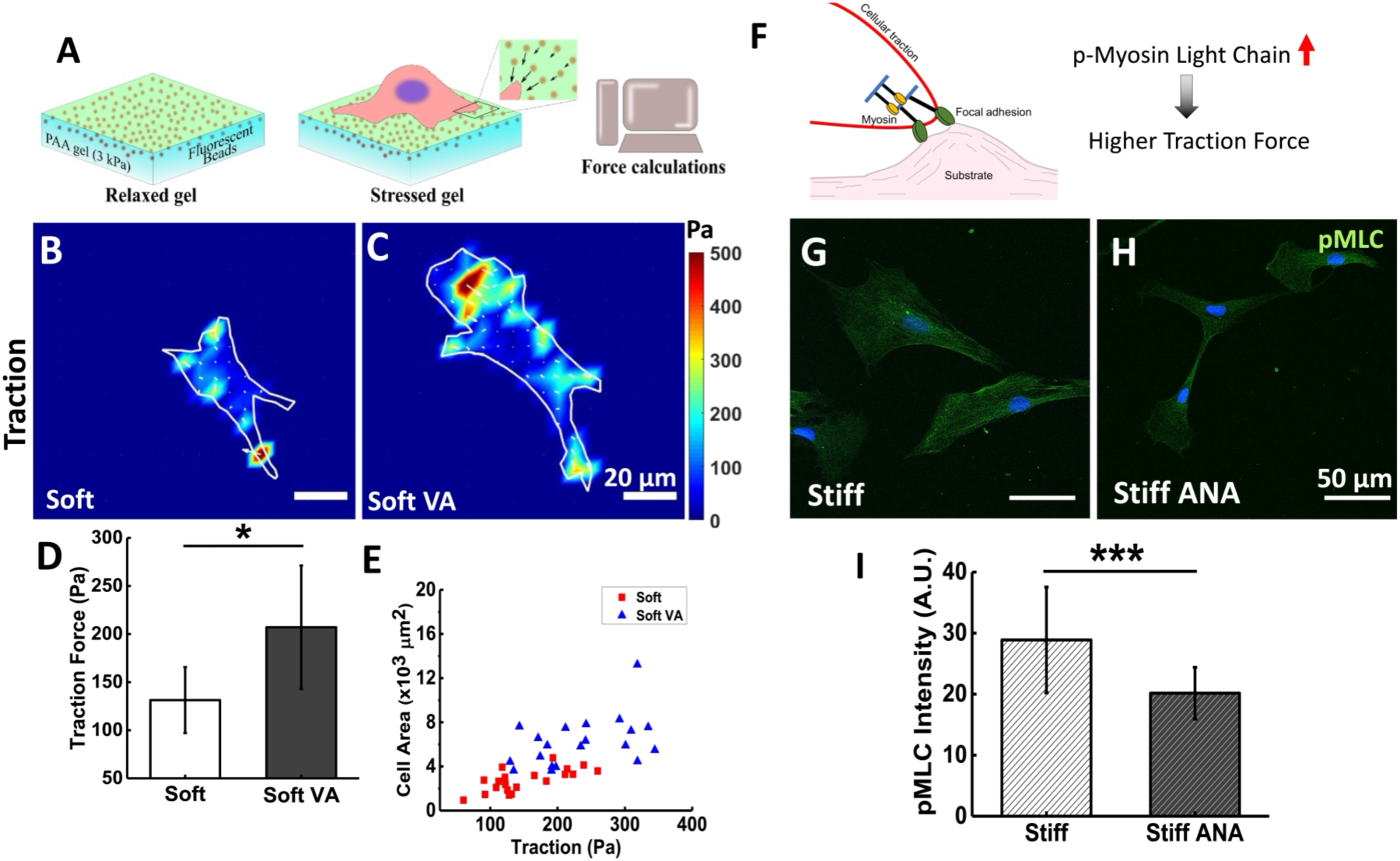
Chromatin remodelling overrides the effect of rigidity on cellular traction of hMSCs. (A) Schematic for PAA hydrogel-based traction force microscopy (TFM) in which cellular traction was estimated from the displacement of the embedded fluorescent beads. (B and C) Heatmap of hMSCs traction force without and with VA. Colours correspond to magnitudes of forces as indicated in the colour bar. (D) Graph shows increase in traction force in addition of VA with respect to control. (E) The schematic representation showing the role of phosphorylated myosin in focal adhesion formation and traction force generation in hMSCs. (F and G) The representative pMLC stained (green) images of the hMSCs cultured on stiff substrate in absence/presence of ANA; scale bar 50 µm. (H) The graph shows the reduction of pMLC intensity upon ANA treatment in hMSCs cultured on stiff substrate.

While TFM provides a robust and quantitative measure of cell-generated forces on soft polyacrylamide (PAA) gels, stiffer substrates (34 kPa) present practical limitations within this experimental framework [29]. As substrate stiffness increases, the deformations induced by cellular forces decrease substantially, leading to bead displacement becoming substantially smaller. This reduction in measurable displacement compromises sensitivity and increases uncertainty in traction reconstruction [30] [31] [32]. Therefore, to assess whether chromatin condensation alters the force generating capacity of hMSCs on stiff substrates, we checked a well-established complementary molecular readout of cellular contractility i.e. phosphorylation of the myosin regulatory light chain (pMLC) (Fig. 3E) [33] [34]. ANA treatment significantly reduced pMLC intensity (from 28.8 ± 8 A.U. to 20.1 ± 4 A.U.) compared to untreated stiff-cultured cells, indicating the reduction in cell’s ability to generate traction force (Figure 3F-H). Collectively, these results demonstrate that chromatin remodeling regulates not only the morphology and cytoskeletal organization but also force generating outputs of hMSCs.

### 4. HDACi/HATi overrides the substrate rigidity-mediated lineage commitment in hMSCs

One of the important characteristics of hMSCs is their multilineage differentiation potential, making them a valuable choice for tissue engineering applications. This lineage commitment in hMSCs is strongly influenced by substrate rigidity and the subsequent changes in cellular contractility, with soft substrate (low contractile state) promoting adipogenic differentiation and stiff substrates (high contractile state) favouring osteogenic differentiation [22], [35], [36], [37], [38]. Since, in our earlier result, chromatin remodelling altered the cellular traction force, we further asked whether it can override these stiffness-dependent lineage cues in hMSCs. Adipogenic and osteogenic differentiation outcomes were assessed following chromatin remodelling by VA and ANA treatment in hMSCs cultured on soft and stiff substrates respectively (Fig. 4A-X).

**Fig. 4.**
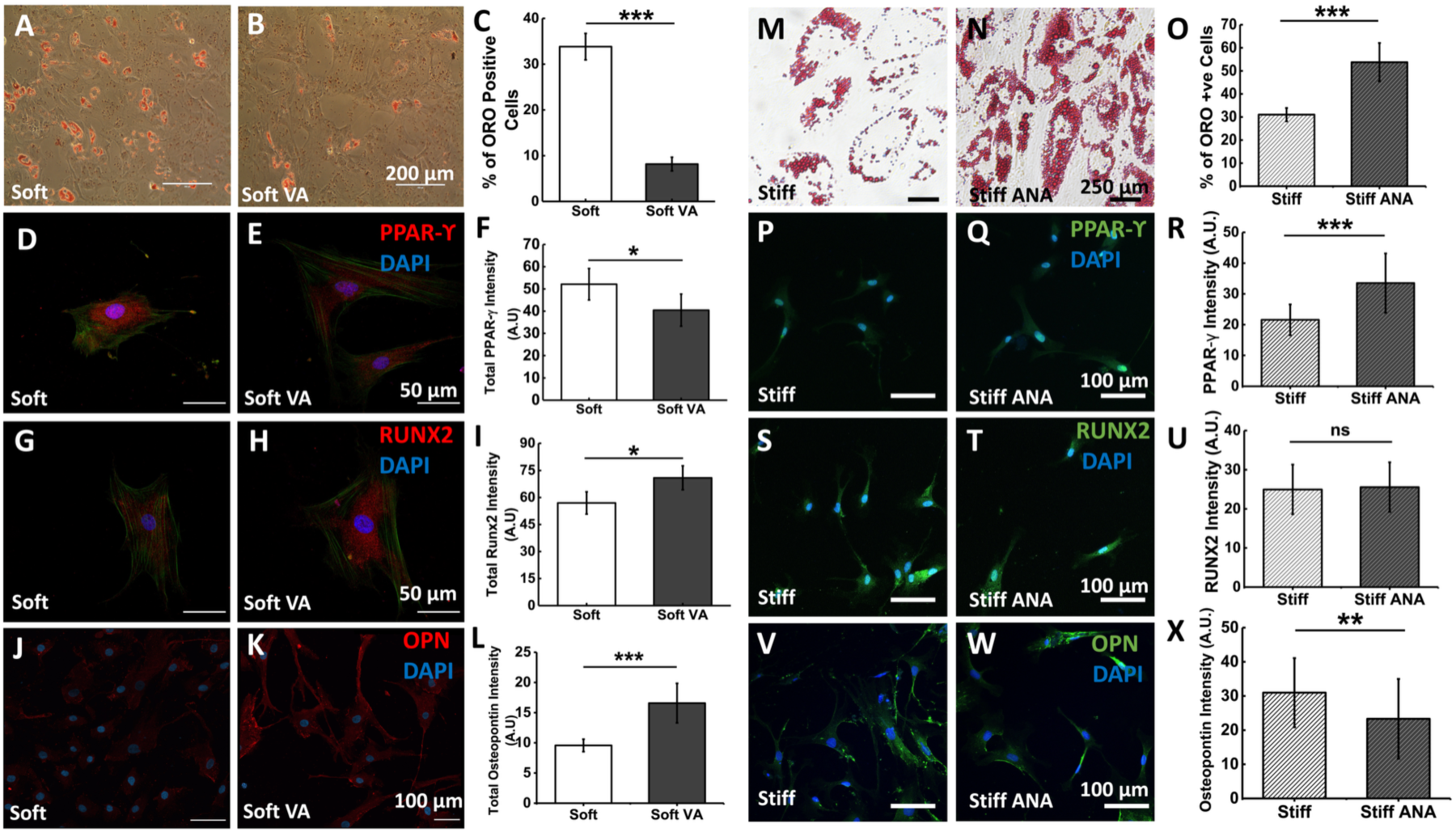
HDACi/HATi overrides the substrate rigidity-mediated lineage commitment in hMSCs: (A and B) The representative images of Oil red O staining after 14 days of culture of hMSCs in adipogenic media on soft gel without and with VA respectively. scale bar: 200 μm (C) The corresponding quantification showing significant drop in adipogenesis upon treatment with VA. (D and E) The adipogenic differentiation has been further demonstrated by staining adipogenic marker PPAR-gamma (red) and DAPI (blue). Scale bar: 50 μm. (F) The PPAR-gamma intensity has been quantified and shown in the graph. (G and H), the representative images of RUNX2 immunostaining (red) on 3 kPa substrate upon inducing osteogenesis in absence and presence of VA. Scale bar 50 μm. (I) The same has been quantified and shown in the graph. (J and K) Another osteogenic marker, Osteopontin (OPN, red) stained cells has been represented when cultured in 3 kPa without and with VA respectively. (L) The OPN intensity has been quantified and shown in the graph. (M and N) The representative images of Oil red O staining after 14 days of culture of hMSCs in adipogenic media on 34 kPa (stiff gel) without and with ANA respectively. Scale bar: 250 μm. (O) The corresponding quantification showing significant increase in adipogenesis upon treatment with ANA. (P and Q) Upon treating ANA to the cells on 34 kPa, the adipogenic marker PPAR-gamma (green) and DAPI (blue) stained cells has been represented. Scale bar: 100 μm. (R) The PPAR-gamma intensity has been quantified and shown in the graph. (S and T) On 34 kPa, the representative images of RUNX2 immunostaining (green) upon inducing osteogenesis in absence and presence of ANA has been shown. Scale bar 100 μm. (U) The RUNX2 intensity in the cells has been quantified and shown in the graph. (V and W) Osteopontin (OPN, green) stained cells has been represented when cultured in 34 kPa without and with ANA respectively. (X) The OPN intensity has been quantified and shown in the graph.

Adipogenic differentiation was assessed by lipid droplet formation and expression of the adipogenic transcription factor PPAR-γ [39], while osteogenic differentiation was evaluated by expression of hallmark proteins such as RUNX2 and osteopontin (OPN) [40], [41]. VA treatment on soft substrates significantly reduced lipid droplet formation by approximately one third compared to untreated controls, reaching levels comparable to those observed on stiff (34 kPa) gels (Fig. 4A–C, S9). Consistent with this reduction in adipogenic commitment, PPAR-γ expression decreased by ∼28% in VA-treated cells relative to untreated control (Fig. 4D-F). A similar reduction in adipogenesis was also observed following treatment with SB (Fig. S10). In contrast, condensing chromatin with ANA treatment significantly increased the fraction of lipid droplet positive cells by ∼1.7 times on stiff substrates compared to untreated controls (Fig. 4M-O). Consistent with this enhanced adipogenic differentiation, PPAR-γ expression was also found to be increased by ∼1.5 times in ANA-treated cells relative to untreated stiff-cultured hMSCs (Fig. 4P-R).

Additionally, upon further increasing chromatin condensation in hMSCs on soft substrate with ANA, we observed a significant reduction in histone acetylation (∼33%; Fig. S14A-C) and increased chromatin condensation (∼23%; Fig. S14D-F) accompanied with increased lipid droplet formation (∼24%) compared to untreated controls (Fig. S14G-I), indicating enhanced adipogenic differentiation.

In parallel, chromatin decondensation promoted osteogenic differentiation on soft substrates. VA treated hMSCs exhibited significantly increased expression of the early osteogenic marker RUNX2 by ∼26% and the late osteogenic marker osteopontin by ∼77% compared to untreated cells cultured on 3 kPa gels (Fig. 4G-L). Quantification of nuclear to cytoplasmic ratios of these markers showed consistent trends, further supporting enhanced osteogenic programming under chromatin-decondensed conditions (Fig. S11).

In contrast, ANA treatment had a differential effect on osteogenic markers. Expression of the early osteogenic transcription factor RUNX2 remained comparable between ANA-treated (25.5 ± 6 A.U.) and untreated cells (24.9 ± 6 A.U.) on stiff substrates (Fig. 4S-U), indicating that chromatin condensation does not impair early osteogenic commitment. However, expression of the late osteogenic marker osteopontin was significantly reduced by ∼1.4 times following ANA treatment (Fig. 4V-X), suggesting that chromatin condensation interferes with osteogenic maturation at later stages.

Collectively, these results demonstrate that chromatin compaction state reprograms stiffness-dependent adipo-osteogenic differentiation in hMSCs. These findings establish chromatin organization as a dominant regulator of lineage specification that can supersede mechanical cues from the ECM.

### 5. Chromatin state overrides rigidity dependent YAP localization in hMSCs

Lineage specification in hMSCs has been linked to the relative nuclear and cytoplasmic localization of Yes-associated protein (YAP), a key mechanosensitive transcription factor which shuttles between the nucleus and cytoplasm in response to matrix stiffness [42] [43]. Given that our results showed chromatin condensation can override substrate stiffness mediated differentiation, where decondensing chromatin on soft substrates promoted osteogenic differentiation and condensing chromatin on stiff substrates promoted adipogenic differentiation; we next examined whether this chromatin-mediated override is accompanied by respective changes in YAP localization. Since YAP nuclear localization is known to be mechanoresponsive, we reasoned that the histone modifier-induced alterations in chromatin state and traction force generation may converge on YAP shuttling as a downstream effector of the observed differentiation outcomes.

To address this question, nuclear and cytoplasmic YAP intensity was measured in hMSCs cultured on soft and stiff gels following chromatin decondensation and condensation respectively (Fig. 5A-L). Decondensing the chromatin with VA in hMSCs cultured on soft substrates increased the nuclear localization of YAP by ∼1.5 times (Fig. 5A-E) and nuclear to cytoplasmic ratio by ∼ 1.4 times (Fig. 5F).

**Fig. 5.**
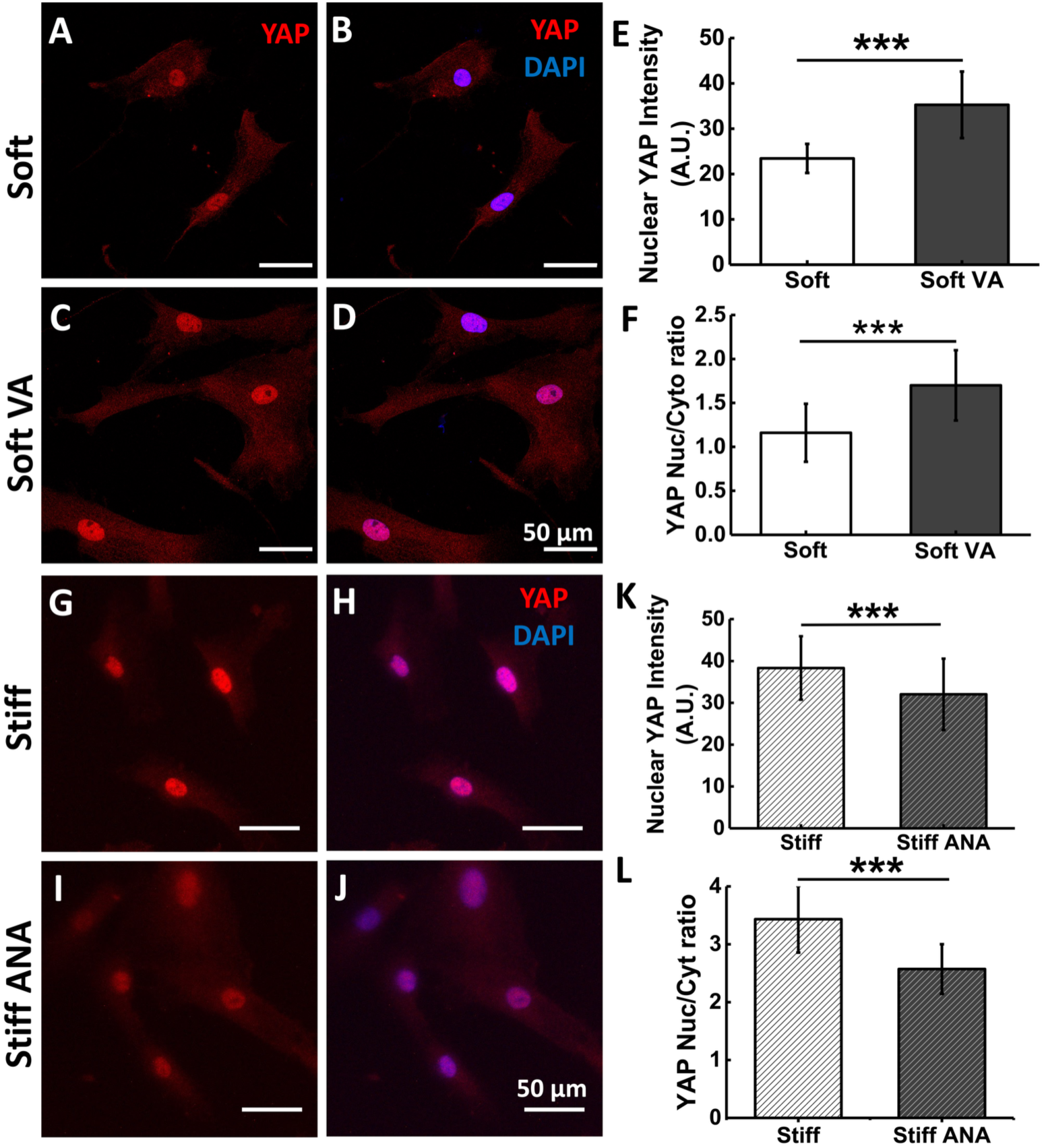
HDACi increases YAP nuclear translocation in hMSCs cultured on soft gel: (A-D) Representative fluorescence images of YAP (red) and nucleus (DAPI: blue) on 3 kPa gel without and with VA. Scale bar: 50 μm. (E and F) The average intensity of nuclear YAP and the ratio of YAP intensity between nucleus and cytoplasm upon VA treatment in hMSCs on soft substrate, (G-J) Representative fluorescence images of YAP (red) and nucleus (DAPI: blue) on 34 kPa gel without and with ANA. Scale bar: 50 μm. (K-L) the average intensity of nuclear YAP and the ratio of YAP intensity between nucleus and cytoplasm upon ANA treatment in hMSCs on stiff substrate.

Similarly, on the stiff substrate condensing chromatin with ANA, decreased the nuclear YAP localization significantly by ∼1.2 times (Fig. 5G-K) and nuclear to cytoplasmic ratio by ∼1.4 times (Fig. 5L) compared to untreated cells, indicating that localization of the YAP was restricted to the cytoplasm. Together, these results demonstrate that chromatin compaction state exerts dominant control over YAP nucleo-cytoplamic localization independent of substrate stiffness.

### 6. ERK Phosphorylation Acts Downstream of Chromatin Compaction to Control Lineage Specification

ERK activity has been reported to be coupled to cellular mechanoresponses, including responses to substrate rigidity, cell spreading, and contractility. Given the central role of ERK/MAPK signalling in coupling extracellular mechanical cues to mesenchymal stem cell fate decisions, we next examined whether chromatin mediated reprogramming of these lineage outcome is accompanied by corresponding changes in ERK activation [44]. Previous studies have shown that ERK phosphorylation is enhanced on stiff substrates, where it promotes osteogenic differentiation, whereas reduced ERK activity on soft substrates favours adipogenesis [45]. Here, we examined whether modulation of chromatin compaction alters ERK signalling independently of substrate stiffness.

To address this, phosphorylated ERK (p-ERK) levels were quantified by immunofluorescence under conditions of chromatin decondensation and condensation. On soft (3 kPa) substrates, treatment with the HDACi resulted in a significant increase in p-ERK intensity (∼35%) compared to untreated controls (Fig. 6A, B, E), indicating activation of ERK signalling despite the soft mechanical environment. In contrast, upon condensing chromatin in hMSCs on stiff (34 kPa) substrates using the HATi (ANA), p-ERK levels were significantly reduced (from 46 ± 9.5 A.U. to 34.9 ± 8 A.U.) compared to untreated cells (Fig. S12). These opposing trends demonstrate that ERK activation closely tracks chromatin state rather than substrate rigidity alone.

**Fig. 6.**
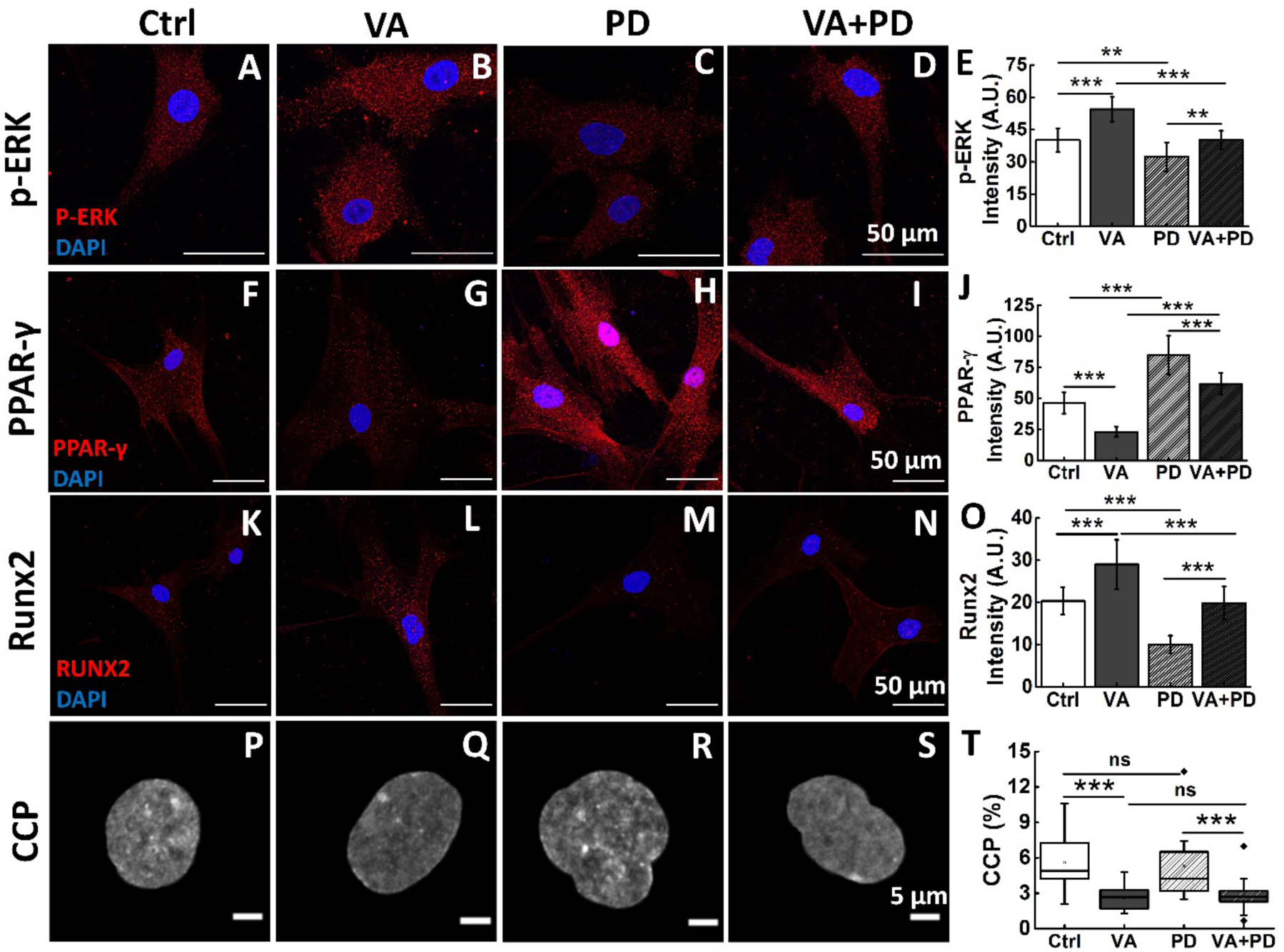
HDACi overwrites the effect of substrate stiffness via p-ERK: On soft substrate, (A–E) p-ERK expression, (F–J) adipogenic marker PPAR-γ expression, and (K–O) osteogenic marker RUNX2 expression were shown in hMSCs treated with VA, PD98059, or VA+PD98059 and compared with untreated controls (CTRL). (P–T) CCP was also measured under the same conditions.

**Fig. 8.**
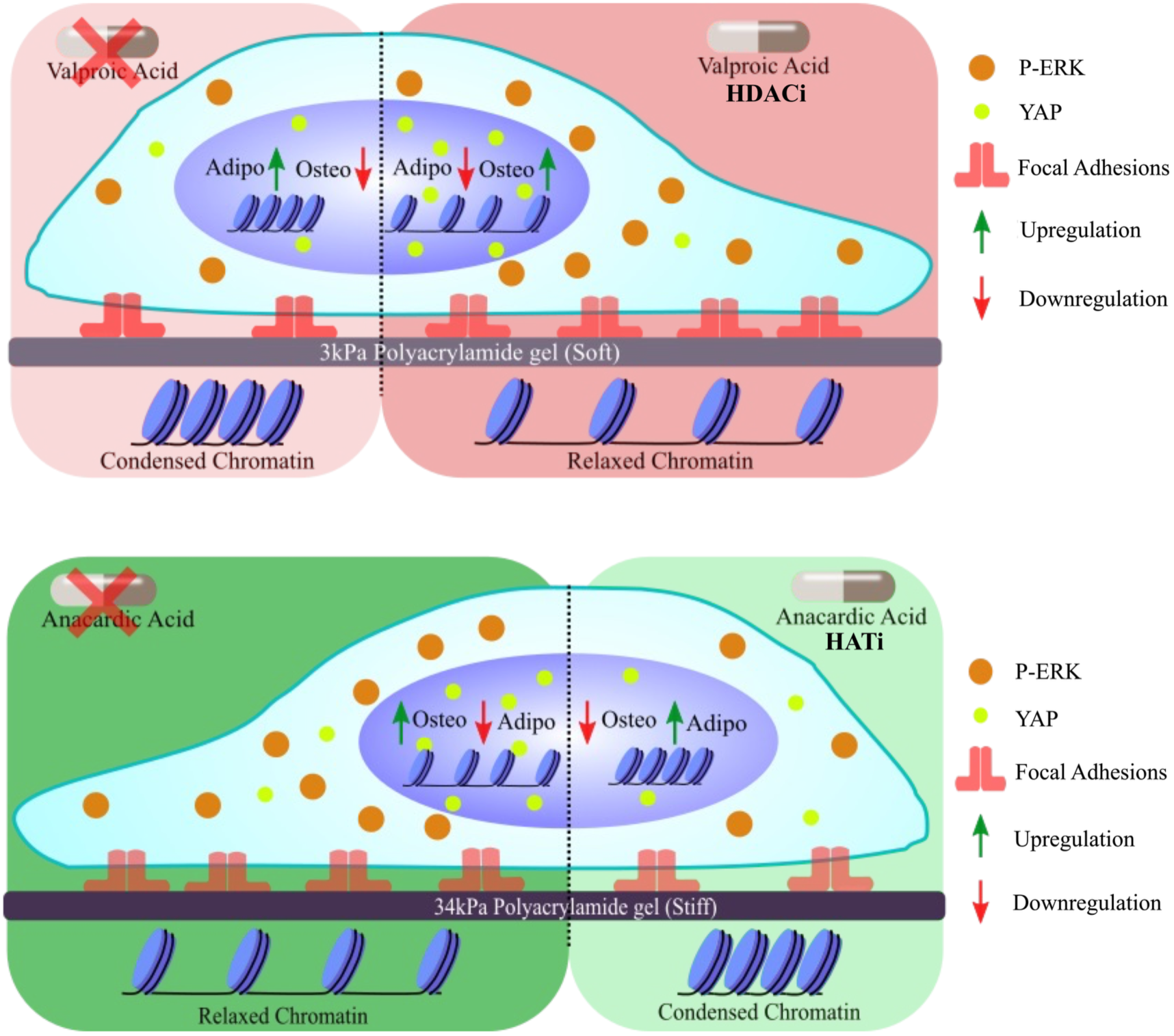
Schematic of summarizing the effect of VA (HDACi) and ANA (HATi) on the mechanoresponse of hMSCs cultured on soft and stiff substrates.

To determine whether ERK activation is functionally required for chromatin driven signalling effects, we inhibited ERK phosphorylation using the ERK inhibitor PD98059 (PD). Treatment of hMSCs with the ERK inhibitor PD98059 (20 µM) resulted in an approximately 25% reduction in p-ERK intensity compared to untreated cells under the same substrate conditions (Fig. 6C-E). Notably, when VA was applied in the presence of PD98059, p-ERK levels were partially restored relative to PD98059 alone (Fig. 6C-E), suggesting that chromatin decondensation can modulate ERK signalling upstream of, or in parallel with, ERK pathway regulation.

We further examined whether p-ERK is required for chromatin compaction-mediated lineage reprogramming by assessing adipogenic and osteogenic marker expression following ERK inhibition under conditions of chromatin decondensation. hMSCs cultured on soft (3 kPa) substrates were treated with VA to induce chromatin decondensation, either alone or in combination with the ERK inhibitor PD98059. Consistent with our earlier observations, VA treatment suppressed adipogenic differentiation on soft substrates, as reflected by reduced PPAR-γ expression (Fig. 6G). Notably, co-treatment with PD98059 restored PPAR-γ expression to levels comparable to control cells (Fig. 6I, J), indicating that inhibition of ERK signalling restored the adipogenic differentiation potential despite chromatin decondensation. As expected, PD98059 treatment alone promoted adipogenesis on soft substrates by blocking ERK phosphorylation (Fig. 6H, J). In parallel, analysis of osteogenic differentiation revealed the opposite trend, VA-induced increase of RUNX2 expression on soft substrates was attenuated upon ERK inhibition, as shown by reduced RUNX2 levels in VA + PD treated cells compared to VA alone (Fig. 6K-O). Importantly, we next asked whether ERK inhibition feeds back to alter chromatin organization itself. Quantification of the chromatin condensation parameter (CCP) showed that VA significantly reduced chromatin condensation (from 5.6 ± 2.26 to 2.6 ± 1.02) relative to untreated controls. However, inhibition of ERK phosphorylation with PD98059 did not affect CCP, either in the presence or absence of VA. These results demonstrate that ERK signalling is not required for chromatin decondensation and instead functions downstream of chromatin state to regulate lineage specification.

Histone acetylation-mediated chromatin remodelling overrides substrate stiffness-dependent mechanoresponse in hMSCs. Soft substrates promote chromatin condensation and adipogenic differentiation, whereas stiff substrates maintain chromatin in a decondensed state and favour osteogenesis. Induction of chromatin decondensation using HDACi drives a stiff-like phenotype on soft substrates, characterized by increased cell spreading, focal adhesion formation, actin stress fibre formation, traction force generation, YAP nuclear localization, and osteogenic differentiation. Conversely, condensing chromatin by HATi promotes a soft-like phenotype in hMSCs on stiff substrates, including reduced spreading, focal adhesion formation, actin expression and enhanced adipogenic differentiation. These findings demonstrate chromatin compaction overrides the substrate rigidity-mediated phenotypes by acting as a central regulator of mechanosignalling and stem cell fate.

## Discussion

hMSCs are the popular choice of stem cells in tissue engineering and regenerative medicines due to their tri-lineage differentiation potential [46], [47]. However, to achieve a desired differentiation outcome, it is important to understand the role of various cues in governing the lineage specification. Matrix stiffness is a dominant biophysical regulator of hMSC behavior and fate, shaping cell spreading, cytoskeletal architecture, traction generation, mechanosensitive signaling, and lineage specification. Although this framework is often presented as a linear pathway, stiffness controls adhesions and contractility, which then drives mechanosensitive cascade. However, it remains unclear how these mechanical cues are integrated at the level of nuclear organization and whether chromatin state can, in turn, reprogram the default stiffness response. In this work, we show that altering histone acetylation dependent chromatin compaction is sufficient to rewire multiple stiffness dependent phenotypes, positioning chromatin compaction also as a central regulatory node for mechanosensitive behavior in hMSCs.

Our results demonstrate that chromatin decondensation induced by VA (HDACi) drives a stiff like cellular program even on soft substrates. VA, an FDA approved anti-cancer drug, has been used in various neurological disorders such as delirium, agitation, epilepsy [48]. Due to its known inhibitory effect in higher concentration (∼4mM) [49], in this work concentration of VA was kept at its minimal level (0.5mM). On soft hydrogels, HDACi promotes enhanced cell spreading, focal adhesion maturation, actin stress-fiber assembly, and higher traction generation; features typically associated with stiff matrices. Conversely, chromatin condensation induced by HATi enforces a soft like program on stiff substrates, reducing spreading, cytoskeletal assembly and adhesion organization despite a mechanically soft environment. This bidirectional control strongly supports a model in which chromatin compaction is not merely correlated with stiffness but can actively determine how cells execute stiffness dependent programs. These results connect to and extend emerging evidence that stiffness influences epigenetic state in hMSCs. For example, it was previously reported that long-term culture and stiffness exposure alter chromatin condensation and histone acetylation, in part through differential expression of HDACs and HATs [7]. Importantly, here in our study we demonstrate that changing chromatin compaction by targeted perturbation of acetylation can supersede the mechanical cue itself, implying that chromatin compaction can function as a dominant integrator of stiffness signals. Conceptually, this positions chromatin not merely as a downstream outcome of mechanosensing, but as an active regulatory element capable of reshaping cell mechanics. Beyond their effects on cellular morphology and mechanophenotype, we found that modulation of chromatin compaction also directly regulates hMSC lineage commitment. While soft substrates typically promote adipogenesis, chromatin decondensation suppressed adipogenic differentiation and enhanced osteogenic markers under these conditions. Conversely, chromatin condensation favoured adipogenic differentiation even on stiff substrates that normally favor osteogenesis. Changes in cytoskeletal organization and YAP localization are often accompanied by alterations in ERK signaling, and ERK levels has been implicated in stiffness dependent mesenchymal stem cell fate decisions. Previous studies have shown that stiff environments show elevated ERK phosphorylation and osteogenic programming, whereas soft substrates are associated with reduced ERK signaling and adipogenic bias. Consistent with these studies, we found that ERK phosphorylation varied with chromatin compaction state across both soft and stiff substrates. If phosphorylation of ERK is inhibited, the chromatin stays in its condensed state but its effect on differentiation disappears and the adipogenic differentiation is restored.

The ability to override stiffness directed fate has practical implications for biomaterials and regenerative medicine. Scaffold mechanics are often tuned to encourage specific differentiation trajectories, yet mechanical design constraints can conflict with optimal stiffness for lineage commitment. A chromatin centered control strategy offers an alternate option for tuning fate outcomes. The findings demonstrated in our study could broaden the range of usable scaffold mechanics by decoupling material stiffness from differentiation bias. Importantly, this approach also provides a framework for understanding how cells resolve conflicting cues, mechanical inputs that bias one lineage can be countered by epigenetic modulation that biases another, and the resulting phenotype reflects the relative dominance of these inputs within a mechanochemical hierarchy.

There are several limitations and opportunities for future work that will further strengthen the mechanistic framework. Future studies that selectively perturb specific HDAC/HAT isoforms, or use CRISPR-based approaches, would help pinpoint which acetylation regulators are most responsible for the observed stiffness override. Second, while we quantify chromatin condensation and signaling outputs, the gene-regulatory mechanisms remain to be mapped. ATAC-seq and targeted ChIP-seq could reveal how chromatin compaction reshapes accessibility of mechanosensitive gene programs and how that connects to ERK-dependent transcriptional states. Finally, extending these experiments to 3D matrices and dynamic mechanical environments will be important, since hMSCs *in vivo* encounter time varying mechanics and complex dimensionality that can reshape both mechanosensing and chromatin state.

In summary, this study demonstrates that chromatin compaction constitutes a key regulatory layer capable of overriding substrate stiffness dependent cues in hMSCs by showing that histone acetylation mediated chromatin remodelling is coupled to coordinated changes in cell adhesion and cytoskeletal organization, traction force generation, YAP localization and lineage specification. Our findings highlight chromatin state as an integrative regulator of mechanosensitive cell behaviour. Together, this work advances a nuclear-centric perspective of mechanoregulation and provides a basis for controlling stem cell fate in mechanically heterogeneous microenvironments.

## Author Information

### Corresponding Author

**Abhijit Majumder**-Department of Chemical Engineering, Indian Institute of Technology Bombay, Mumbai, Maharashtra, India-400076.

**Rohit Joshi**-Department of Chemical Engineering, Indian Institute of Technology Bombay, Mumbai, Maharashtra, India-400076.

### Authors

**Sourav Mukherjee** - Department of Chemical Engineering, Indian Institute of Technology Bombay, Mumbai, Maharashtra, India-400076.

**Pooja Dharmambal Muralidharan-** Department of Chemical Engineering, Indian Institute of Technology Bombay, Mumbai, Maharashtra, India-400076.

**Tejas Suryawanshi** - Center for Research in Nanotechnology and Science, Indian Institute of Technology Bombay, Mumbai, Maharashtra, India-400076.

**Pushpendra Yadav-** Department of Chemical Engineering, Indian Institute of Technology Bombay, Mumbai, Maharashtra, India-400076.

**Vedanshi Dharnidharka-** Department of Chemical Engineering, Indian Institute of Technology Bombay, Mumbai, Maharashtra, India-400076.

**Prof. Shobha Shukla** - Department of Metallurgical Engineering and Materials Science, Indian Institute of Technology Bombay, Mumbai, Maharashtra, India-400076.

## Author Contribution

A.M., R.J. and S.M. conceptualized and designed the research; A.M. supervised the project. R.J., S.M., P.M., T.S., P.Y. and V.D. performed the experiments. R.J., S.M., T.S., P.M., P.Y. and V.D. analyzed the data. R.J, S.M., prepared the figures. R.J., S.M. and A.M., S.S. interpreted the results. R.J., S.M., T.S., A.M. drafted the manuscript.

## Data Availability

All relevant data and associated protocols are provided within the article and *SI Appendix*.

## Acknowledgement

AM acknowledge ANRF/ARG/2025/003146/LS and CRG/2022/005992 for funding support. RJ, SM acknowledge DBT/2016/IIT-B/560, Industrial Research And Consultancy Centre (IRCC, IIT Bombay) and 17X(3)/Adhoc/65/2022-ITR for providing fellowship support. We thank the confocal microscopy facility of IIT Bombay. We thank Dr. Subhajit Sen for discussion and providing sodium butyrate. We thank Dr. Akshada Khadpekar for helping in running CCP MATLAB script. We thank Dr. James P Butler (Harvard Medical School, Department of Medicine, Boston) for his TFM code used for the analysis.

## SUPPLEMENTARY INFORMATION

**Fig. S1.**
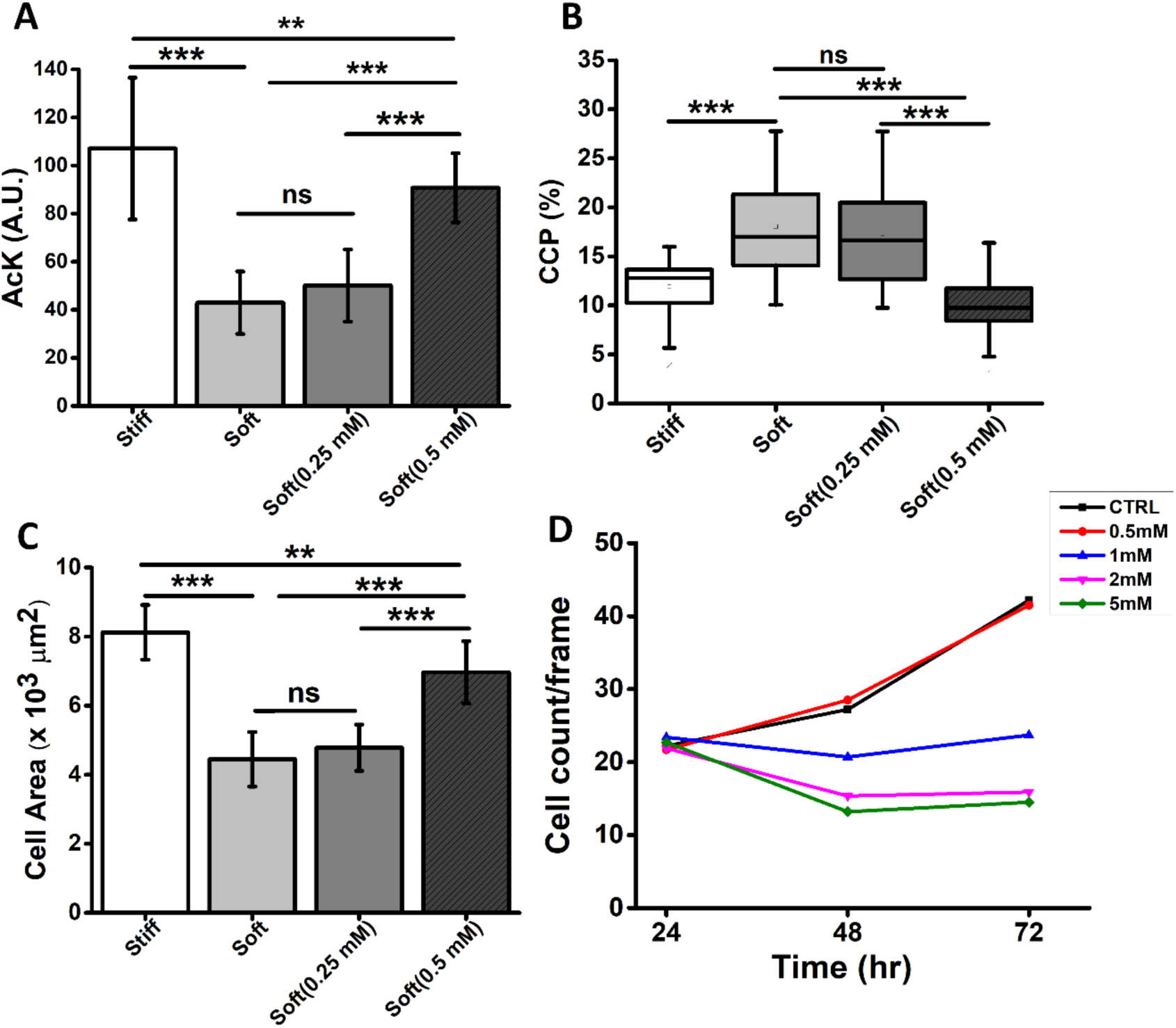
Effect of substrate stiffness and varying concentration of VA (HDACi) on spread area, acetylation and CCP. Quantification showing change in (A) Acetylation (B) chromatin condensation parameter (C) cell spread area. Graph shows significant change in cell area, AcK and CCP at 0.5 mM but no change at lower concentration (0.25 mM). Graph (D) shows at 0.5 mM of HDACi there is no effect on cell growth as compared to control. *p<0.05, **p<0.01, ***p<0.001, N=3,n=50.

**Fig. S2.**
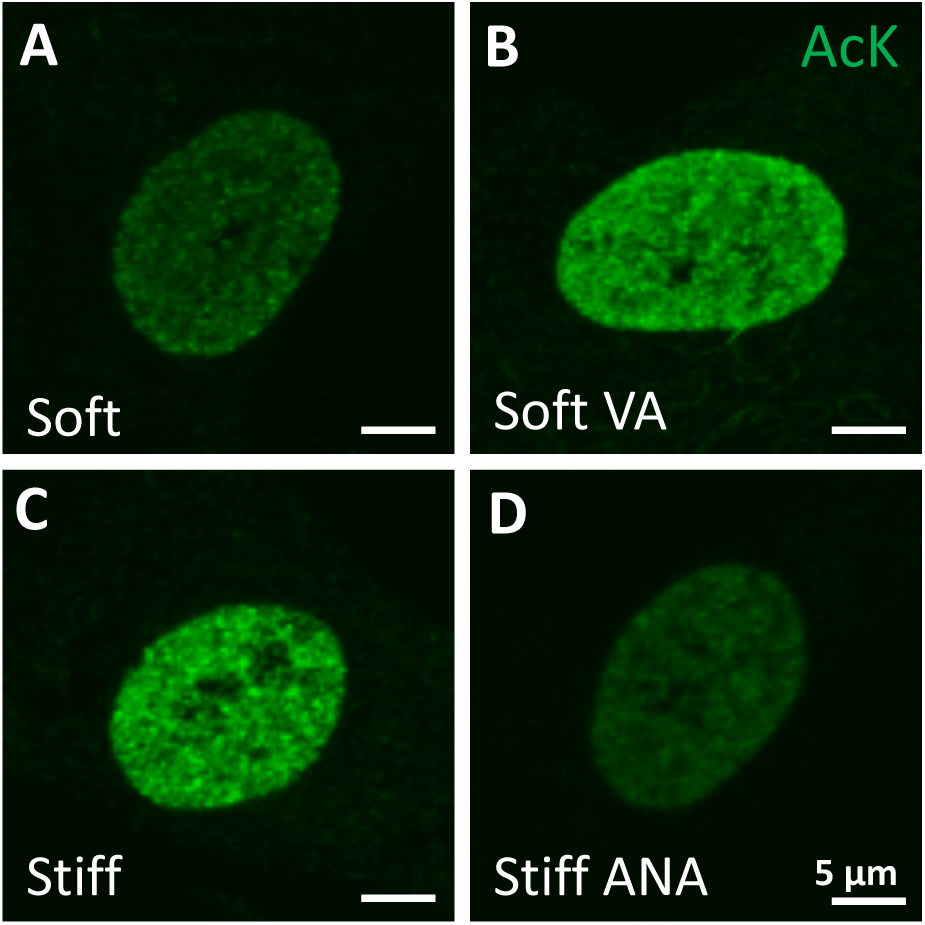
Influence of substrate stiffness and HDACi/HATi on chromatin condensation. (A-B) Representative zoomed-in immunostained images showing intensity of acetylation of lysin residue in histones shown in green for cells cultured on soft (A), soft + VA (B), stiff (C) and stiff +ANA (D); scale bar: 5 µm.

**Fig. S3.**
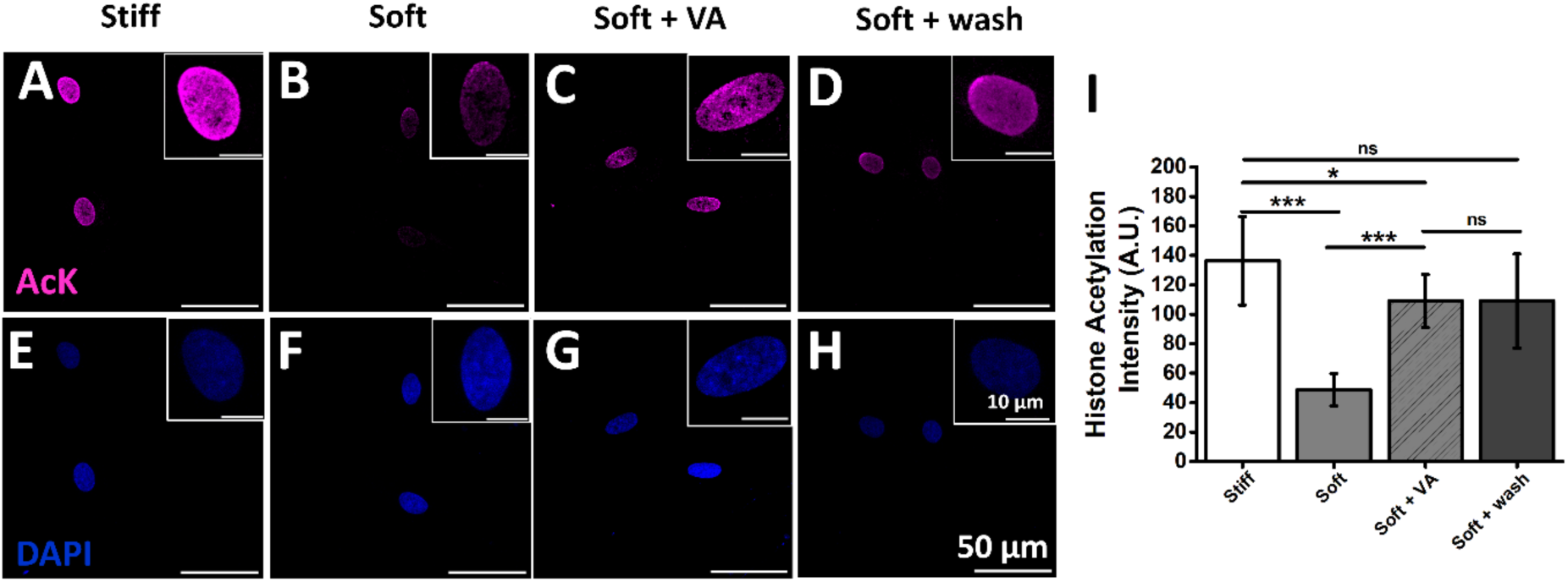
Effect of HDACi washing on histone acetylation: (A-H) Representative immunostained images showing average intensity of acetylation of lysine residue in histones shown in magenta and respective nucleus (DAPI-blue) for cells cultured on stiff (34 kPa, A and E), soft (3 kPa, B and F), soft + VA (C and G) and soft + wash (D and H). Quantification of the same is shown in Fig. I. Our result shows that even after 24 h of washing of VA, the chromatin remains in the open state. *p<0.05, **p<0.01, ***p<0.001, N=3, n>30. scale bar: 50 µm (H) & inset: 10 µm (H).

**Fig. S4.**
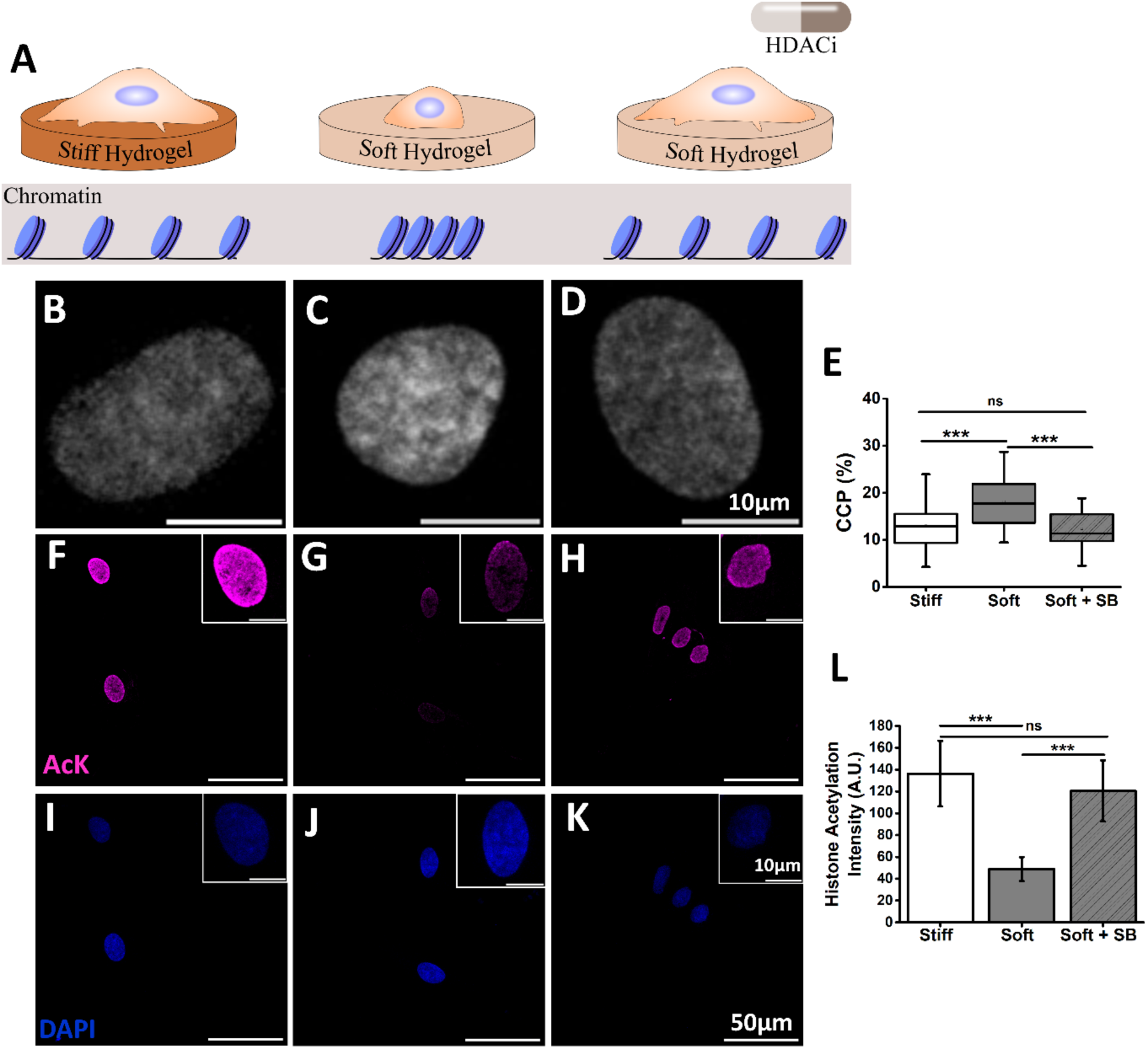
Influence of substrate stiffness and sodium butyrate (HDACi) on chromatin condensation. (A) A schematic showing changes in chromatin condensation due to substrate stiffness and HDACi (sodium butyrate, SB). (B-D) Representative images of DAPI-stained nuclei from stiff (34 kPa), soft (3 kPa) and soft + SB. (E) Quantification of CCP defined as % of highly DAPI stained regions to the total nuclear area shown for nuclei from stiff, soft and soft + SB. The result demonstrates that while soft substrate increases chromatin condensation, the same can be brought down to the level of stiff substrates by treating the cells with SB. (F-K) Representative immunostained images showing average intensity of acetylation of lysine residue in histones shown in magenta and respective nucleus (DAPI-blue) for cells cultured on stiff (34 kPa, F and I), soft (3 kPa, G and J) and soft + SB (H and K). Quantification of the same is shown in Fig. L. *p<0.05, **p<0.01, ***p<0.001, N=3, n>30. scale bar: 10 µm (D), 50 µm (K) & inset: 10 µm (K).

**Fig. S5.**
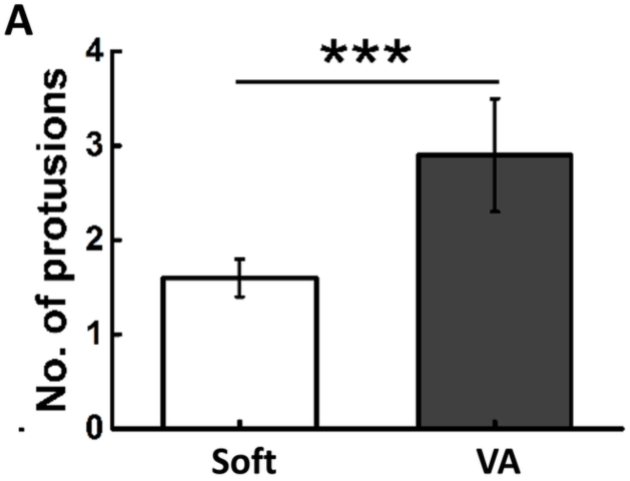
Effect of HDACi on focal adhesion area and actin anisotropy. Graph (A) show quantification of no. of protrusions in each cell when cultured on soft gel with and without VA. *p<0.05, **p<0.01, ***p<0.001.

**Fig. S6.**
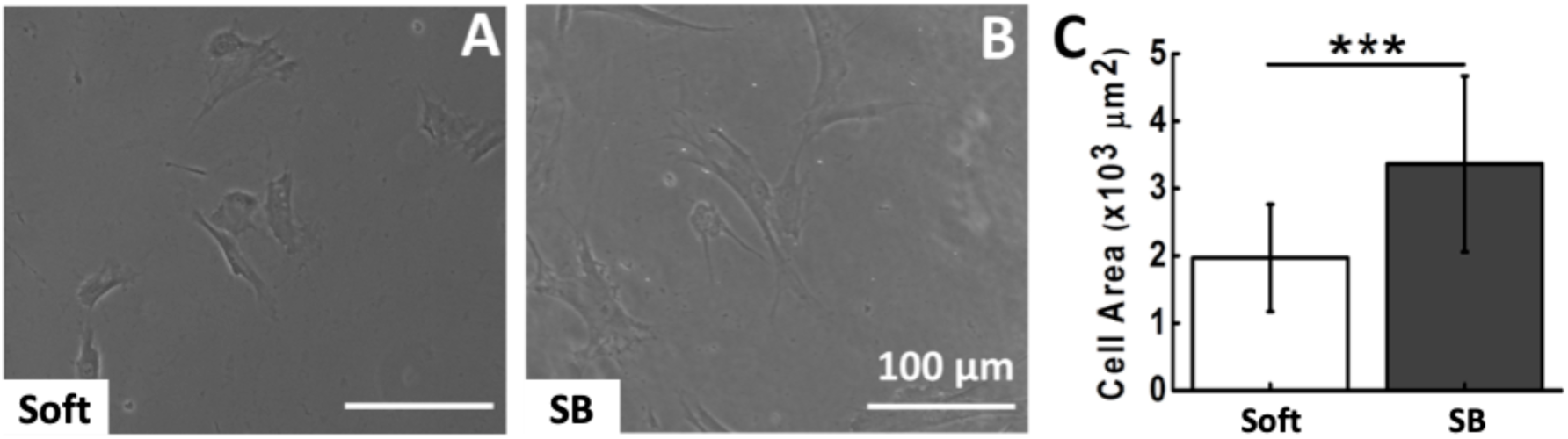
HDACi increases cell spread area on soft substrate. Representative images of hMSCs cultured on 3 kPa gel (A) without SB and (B) with SB. Graph (C) show quantitative analysis of change in projected cell area on 3 kPa gels with and without SB. \**p*<0.05, **p<0.01, ***p<0.001, N=3, n > 50 cells, scale bar: 100 μm.

**Fig. S7.**
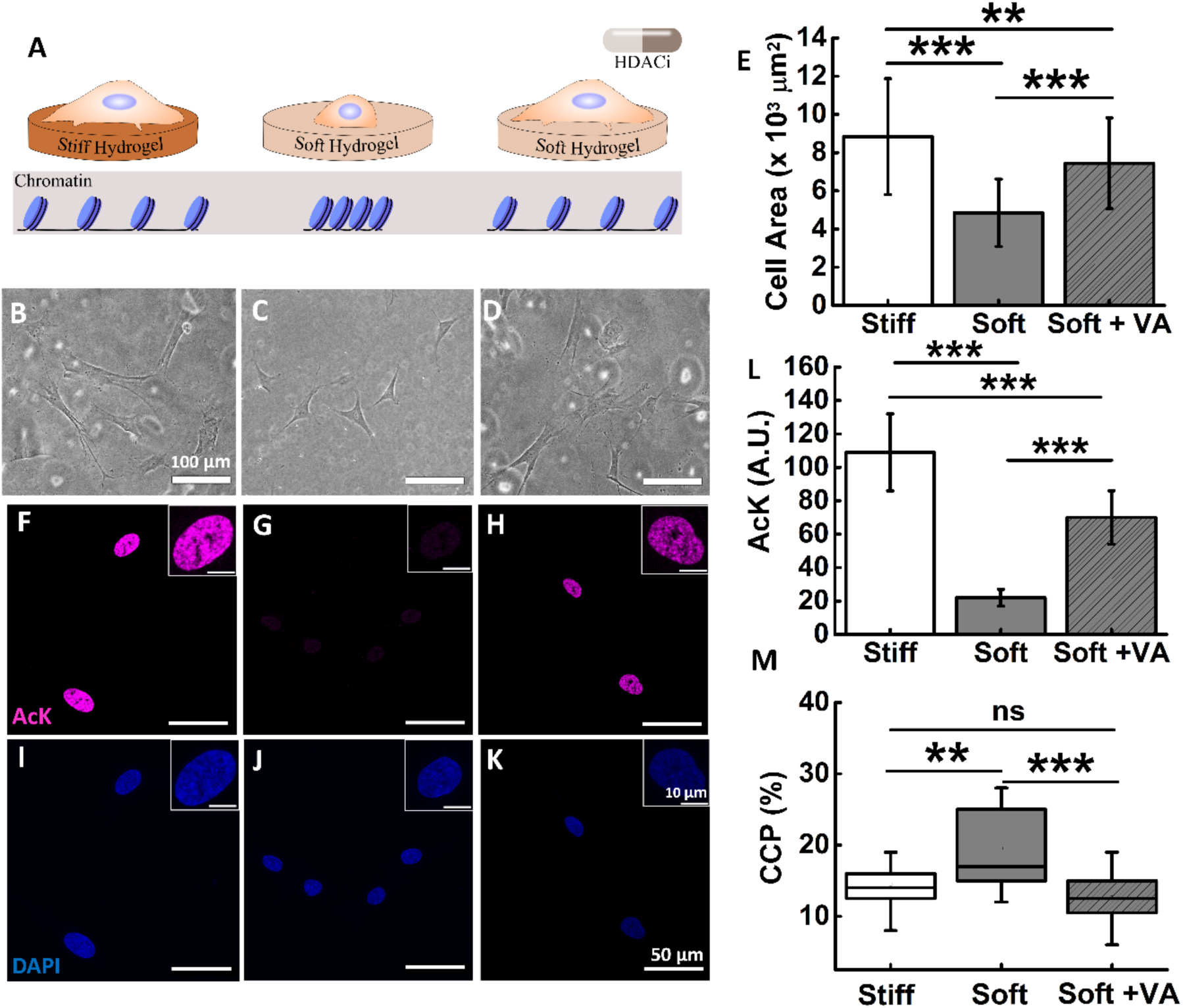
Influence of fibronectin coating on hMSCs morphology and chromatin condensation in presence of HDACi on stiff and soft substrates. (A) schematic showing changes in cellular morphology and chromatin condensation due to substrate stiffness and HDACi (VA). (B-D). Representative phase contrast images of hMSCs cultured on stiff (34 kPa), soft (3 kPa) and soft + VA. (F-H) Representative images of acetylation (magenta) and (I-K) respective nucleus (DAPI-blue) for cells cultured on stiff (34 kPa), soft (3 kPa) and soft + VA. Graph (E) shows quantification of cell spread area of hMSCs cultured on stiff (34 kPa), soft (3 kPa) and soft + VA. Graph ( L & M) shows quantification of average acetylation intensity and CCP of cells cultured on stiff (34 kPa), soft (3 kPa) and soft + VA. *p<0.05, **p<0.01, ***p<0.001, N=3, n>50, scale bar: 100 μm (B), 50 μm (K), inset : 10µm (K).

**Fig. S8.**
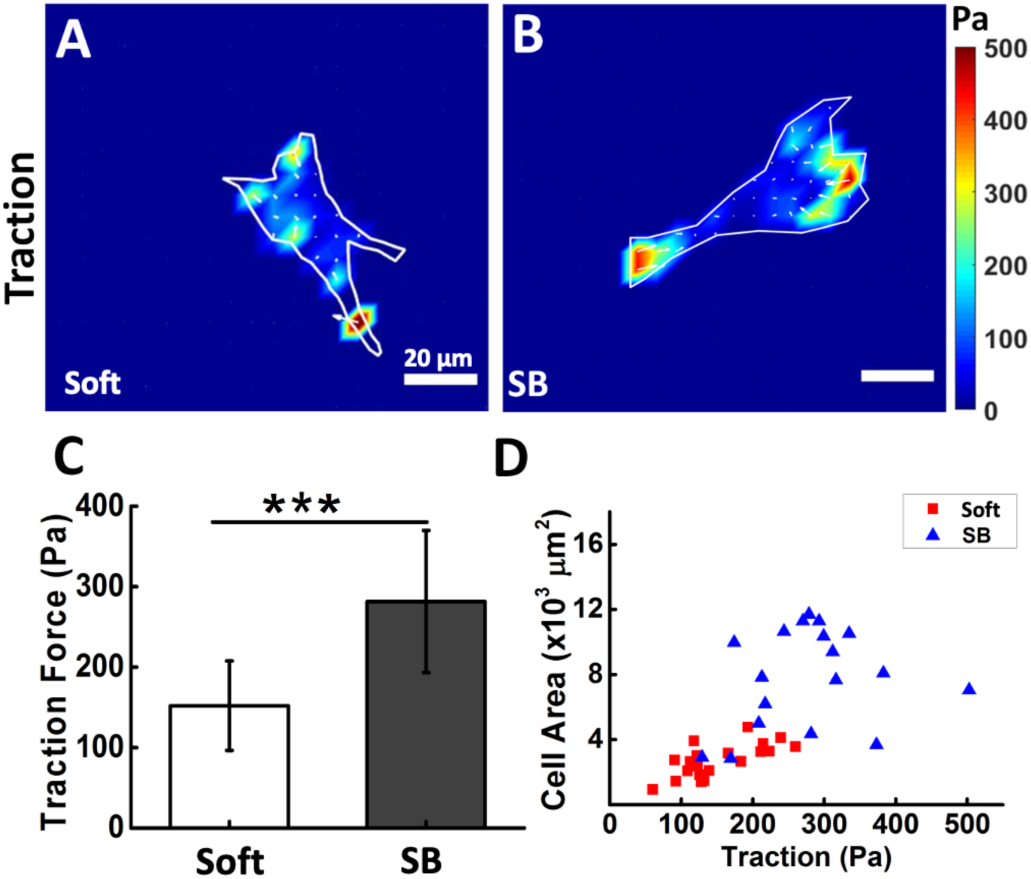
HDACi, sodium butyrate increases hMSCs traction on soft substrates. Heat map of soft substrate-cultured hMSCs traction stress (A) without SB, (B) with SB. Colours corresponds to magnitude of stresses as indicated in the colour bar. Graph (C) shows quantitative analysis of traction force on 3 kPa gel with and without SB (Ctrl). Scatter plot (D) shows positive correlation between cell spread area and cellular traction. \**p*<0.05, **p<0.01, ***p<0.001, N=2, n>15 cells, scale bar: 20 μm.

**Fig. S9.**
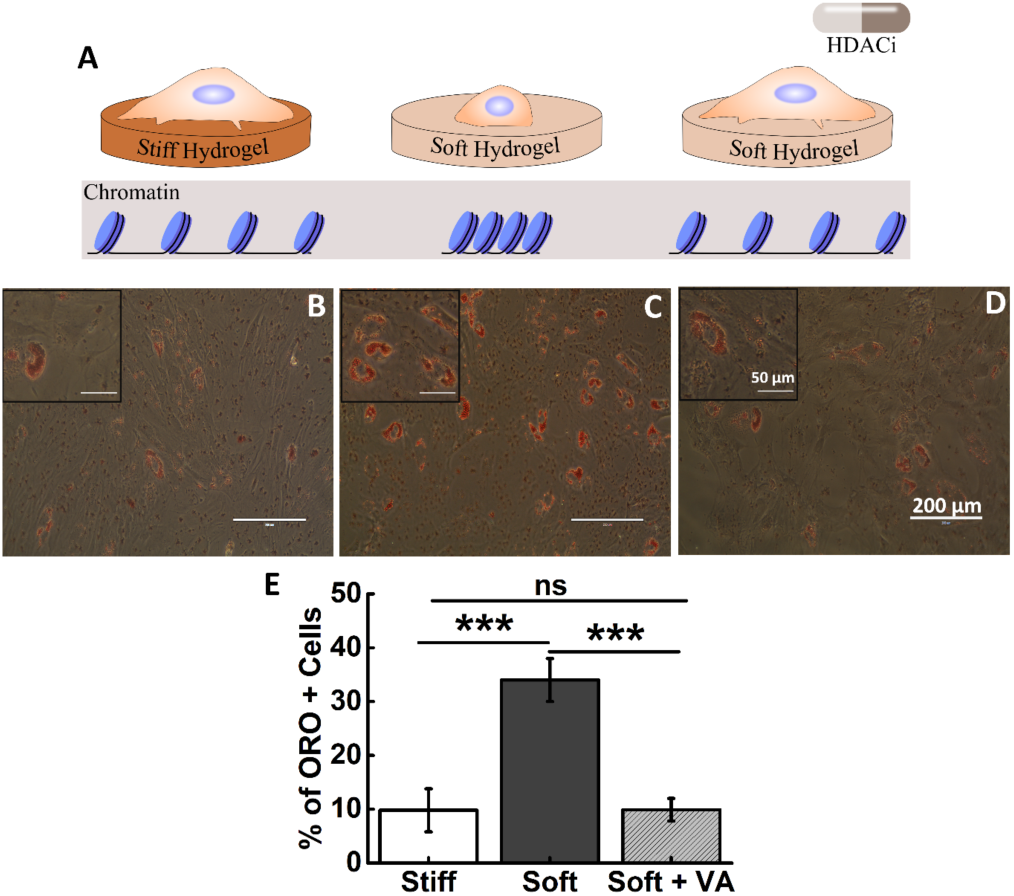
HDACi overrides the adipogenic differentiation potential of hMSCs cultured on soft substrates (3 kPa). (A) schematic showing changes in chromatin condensation due to substrate stiffness and HDACi. Representative images of Oil Red O staining for hMSCs cultured on stiff (34 kPa), soft (3 kPa) and soft + VA. The insets are the magnified local image to show Oil Red O formation. Graph (E) shows quantitative analysis of change in percentage of ORO positive cells for hMSCs cultured on stiff (34 kPa), soft (3 kPa) and soft + VA. *p<0.05, **p<0.01, ***p<0.001, N=3, n=300, scale bar: 200 μm, inset : 50 µm.

**Fig. S10.**
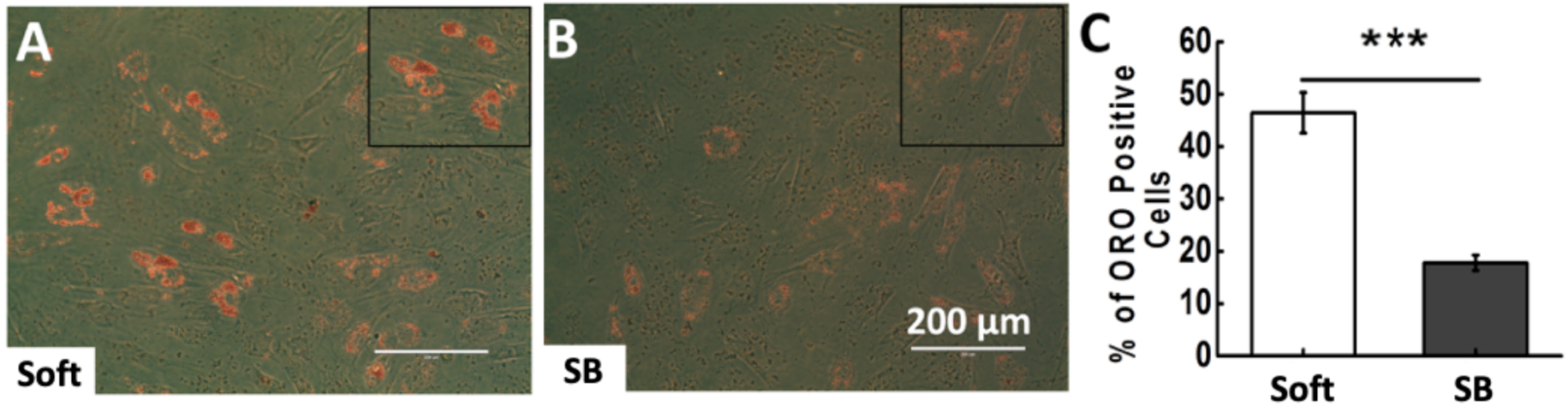
HDACi inhibit adipogenesis of hMSCs cultured on soft substrates. Representative images of Oil Red O staining on 3 kPa gel without SB (A) and with SB (B). The insets are the magnified local image to show oil droplets. Graph (C) shows quantitative analysis of change in percentage of ORO positive cells with and without SB. \**p*<0.05, **p<0.01, ***p<0.001, N=3, n=300, scale bar: 200 μm.

**Fig. S11.**
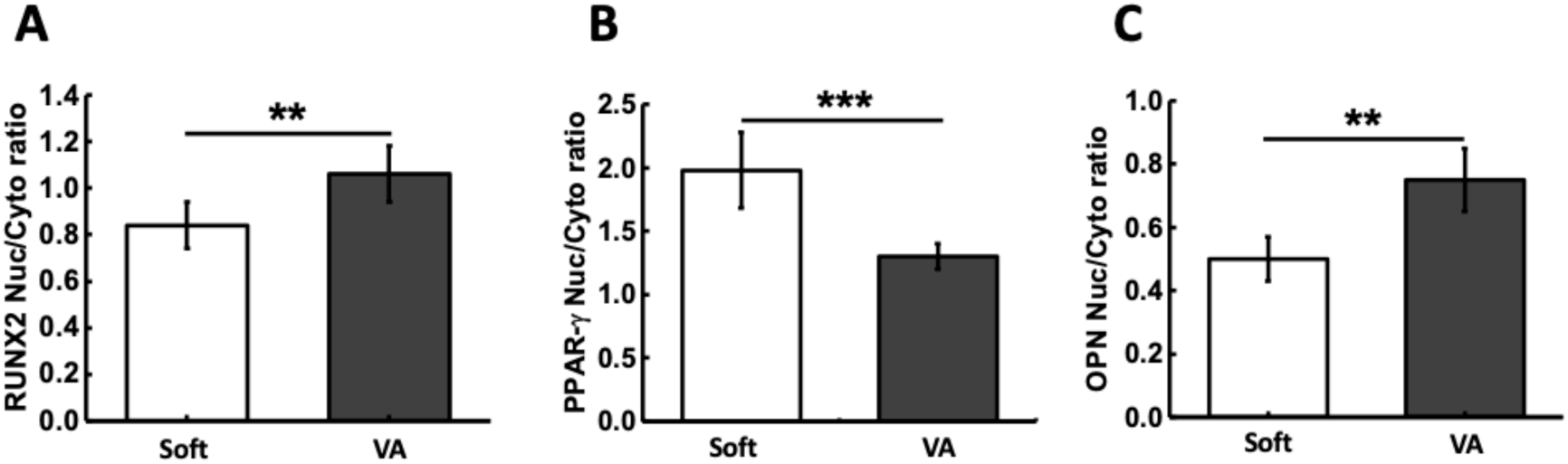
Effect of HDACi on nuclear/cytoplasmic ratio of differentiation markers. Graph (A), (B) and (C) show quantification of RUNX2, PPAR-γ and osteopontin nuclear to cytoplasmic ratio respectively, when the cells are cultured on soft gel with and without VA. *p<0.05, **p<0.01, ***p<0.001, N=3, n>25.

**Fig. S12.**
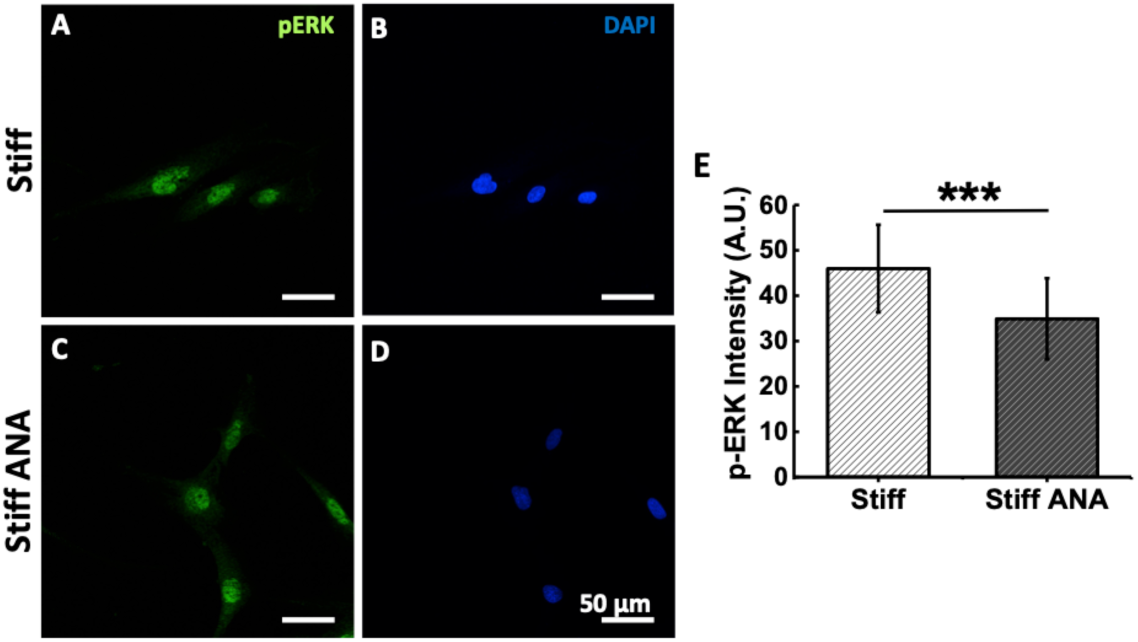
Effect of HATi on pERK expression on stiff substrate. (A-D) Representative p-ERK (green) and DAPI (blue) stained immunofluorescence images has been shown for hMSCs cultured on 34 kPa substrate without and with ANA respectively. (G) The graph of quantitative measurement of p-ERK intensity in hMSCs has been shown. *p<0.05, **p<0.01, ***p<0.001, N=3, n>25.

**Fig. S13.**
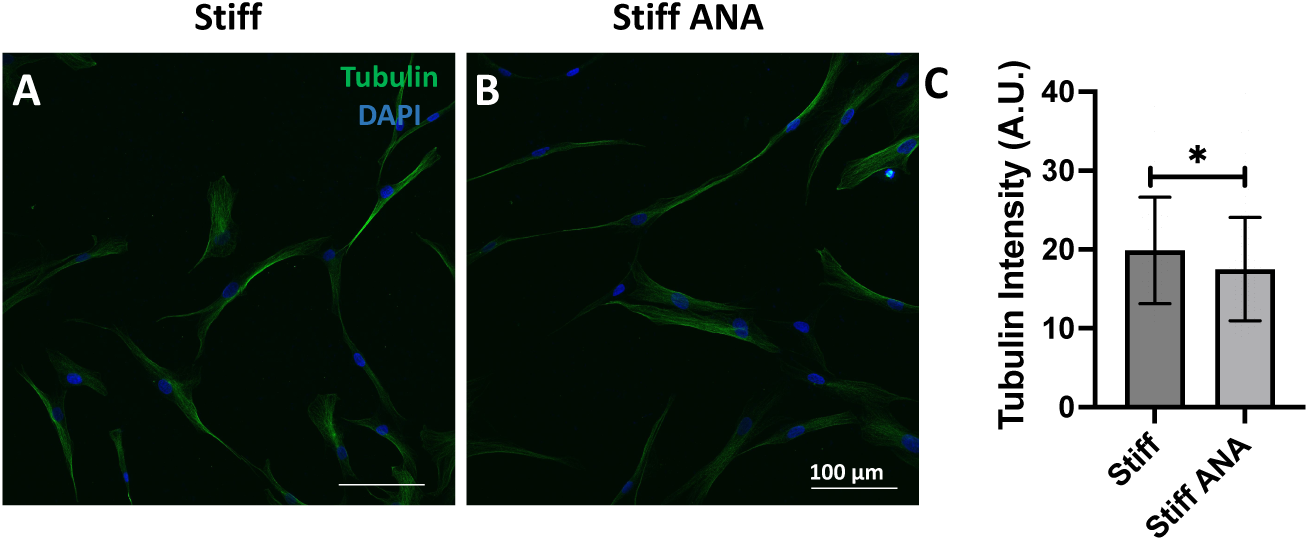
Effect of HATi on Tubulin expression on stiff substrate. (A and B) Representative immunofluorescence images of Tubulin (green) and DAPI (blue), along with the corresponding merged panels, for hMSCs cultured on 34 kPa substrates without and with ANA, respectively. (C) The quantitative measurement of Tubulin intensity in hMSCs has been shown. *p<0.05, n>50.

**Fig. S14.**
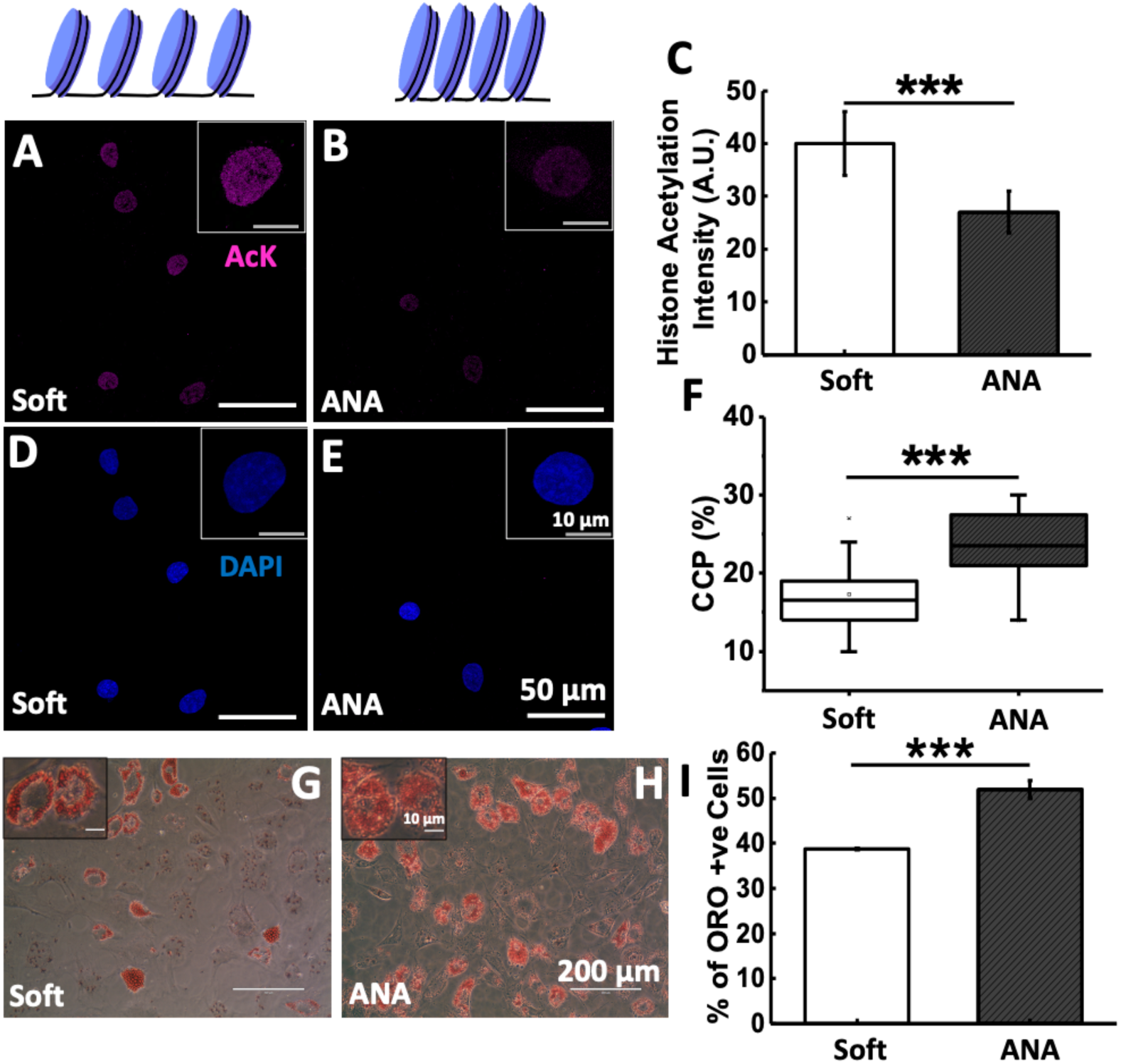
Chromatin condensation promotes adipogenesis of hMSCs cultured on soft substrate: (A-B) Representative images of acetylation (magenta) and (D-E) respective nucleus (DAPI-blue) from Soft (3 kPa) (A and D) and Soft + ANA (B and E). (C) shows corresponding quantification of Histone acetylation average intensity. (F) Shows quantification of % CCP for the nuclei from Soft (3 kPa) and Soft + ANA. Oil Red O staining after ten days of culturing hMSCs in adipogenic induction media (AIM) on soft gel (3 kPa) without (G) and (H) with ANA. (I) Quantification of the percentage of lipid oil droplets in the presence of AIM and AIM with ANA shows that treatment with ANA further increases adipogenesis on soft gel. *p<0.05, **p<0.01, ***p<0.001, N=3, n>30. Scale bar:50 µm (E), inset:10 µm (E), 200 µm (H), inset:10 µm (H).

**Table S1:** Polyacrylamide Gel composition of soft (3 kPa) and stiff (34 kPa) gels.

| Acrylamide from<br>40% stock solution (mL) | Bis-acrylamide from<br>2% Stock solution (mL) | Water<br>(mL) | E ± St. Dev.<br>(kPa) |
| --- | --- | --- | --- |
| 1 | 1.125 | 7.875 | 3.13 ± 0.42 |
| 2.5 | 1.5 | 6 | 34.88 |

## References

[1] M. Ehrbar et al., “Elucidating the Role of Matrix Stiffness in 3D Cell Migration and Remodeling,” Biophys. J., vol. 100, no. 2, pp. 284–293, Jan. 2011, doi: 10.1016/j.bpj.2010.11.082.

[2] B. L. Doss et al., “Cell response to substrate rigidity is regulated by active and passive cytoskeletal stress,” Proc. Natl. Acad. Sci. U. S. A., vol. 117, no. 23, pp. 12817–12825, Jun. 2020, doi: 10.1073/pnas.1917555117.

[3] J. Morival, A. Hazelwood, and J. Lammerding, “Feeling the force from within – new tools and insights into nuclear mechanotransduction,” J. Cell Sci., vol. 138, no. 5, p. JCS263615, Mar. 2025, doi: 10.1242/jcs.263615.

[4] Y. Zhan, S. Shao, B. Liu, and X. Yu, “Stress transmission towards the nucleus of the cell,” Front. Cell Dev. Biol., vol. 14, Mar. 2026, doi: 10.3389/fcell.2026.1790136.

[5] P. S. Mathieu and E. G. Loboa, “Cytoskeletal and Focal Adhesion Influences on Mesenchymal Stem Cell Shape, Mechanical Properties, and Differentiation Down Osteogenic, Adipogenic, and Chondrogenic Pathways,” Tissue Eng. Part B Rev., vol. 18, no. 6, p. 436, Dec. 2012, doi: 10.1089/TEN.TEB.2012.0014.

[6] R. Joshi, D. Shah, K. V. Venkatesh, and A. Majumder, “Histone acetylation alters the nuclear morphology and architecture of human mesenchymal stem cells in a rigidity-dependent manner,” Soft Matter, vol. 21, no. 48, pp. 9230–9244, Dec. 2025, doi: 10.1039/d5sm00941c.

[7] A. R. Killaars, J. C. Grim, C. J. Walker, E. A. Hushka, T. E. Brown, and K. S. Anseth, “Extended Exposure to Stiff Microenvironments Leads to Persistent Chromatin Remodeling in Human Mesenchymal Stem Cells,” Adv. Sci., vol. 6, no. 3, p. 1801483, 2019, doi: 10.1002/advs.201801483.

[8] J. Buisson, X. Zhang, T. Zambelli, P. Lavalle, D. Vautier, and M. Rabineau, “Reverse Mechanotransduction: Driving Chromatin Compaction to Decompaction Increases Cell Adhesion Strength and Contractility,” Nano Lett., vol. 24, no. 14, pp. 4279–4290, Apr. 2024, doi: 10.1021/acs.nanolett.4c00732.

[9] “Chromatin enables precise and scalable gene regulation with factors of limited specificity | PNAS.” Accessed: Sep. 11, 2026. [Online]. Available: https://www.pnas.org/doi/10.1073/pnas.2411887121

[10] A. D. Stephens, “Chromatin Rigidity Provides Mechanical and Genome Protection,” Mutat. Res., vol. 821, p. 111712, 2020, doi: 10.1016/j.mrfmmm.2020.111712.

[11] M. Rabineau et al., “Chromatin de-condensation by switching substrate elasticity,” Sci. Rep., vol. 8, no. 1, p. 12655, Aug. 2018, doi: 10.1038/s41598-018-31023-2.

[12] Suarez Ivan Rey, A. Pathni, F. Fazekas, M. Connell, A. Upadhyaya, and T. Misteli, “Nuclear morphology and chromatin compaction modulate T cell cytoskeletal remodeling and immune synapse formation,” Mol. Biol. Cell, vol. 37, no. 9, p. ar83, Sep. 2026, doi: 10.1091/mbc.E25-10-0487.

[13] A. R. Killaars, J. C. Grim, C. J. Walker, E. A. Hushka, T. E. Brown, and K. S. Anseth, “Extended Exposure to Stiff Microenvironments Leads to Persistent Chromatin Remodeling in Human Mesenchymal Stem Cells,” Adv. Sci., vol. 6, no. 3, p. 1801483, 2019, doi: 10.1002/advs.201801483.

[14] H. Romero, et al., “Chromatin organization controls nuclear stiffness,” Mar. 15, 2025, bioRxiv. doi: 10.1101/2025.03.14.643219.

[15] R. Karki, F. N. D. L. Vitorino, R. M. Searfoss, J. K. Lempiäinen, C. H. Arrowsmith, and B. A. Garcia, “Comprehensive mass spectrometry screening-derived atlas of HDAC inhibitors reveals histone-specific acetylation changes,” Cell Rep., vol. 45, no. 5, p. 117289, May 2026, doi: 10.1016/j.celrep.2026.117289.

[16] M. Huang, J. Huang, Y. Zheng, and Q. Sun, “Histone acetyltransferase inhibitors: An overview in synthesis, structure-activity relationship and molecular mechanism,” Eur. J. Med. Chem., vol. 178, pp. 259–286, Sep. 2019, doi: 10.1016/j.ejmech.2019.05.078.

[17] J. R. Tse and A. J. Engler, “Preparation of Hydrogel Substrates with Tunable Mechanical Properties,” Curr. Protoc. Cell Biol., vol. 47, no. 1, p. 10.16.1-10.16.16, 2010, doi: 10.1002/0471143030.cb1016s47.

[18] R. Vishavkarma, S. Raghavan, C. Kuyyamudi, A. Majumder, and J. Dhawan, “Role of Actin Filaments in Correlating Nuclear Shape and Cell Spreading,” PLoS ONE, vol. 9, no. 9, p. 107895, 2014, doi: 10.1371/journal.pone.0107895.

[19] J. Irianto et al., “Osmotic Challenge Drives Rapid and Reversible Chromatin Condensation in Chondrocytes,” Biophys. J., vol. 104, no. 4, pp. 759–769, Feb. 2013, doi: 10.1016/j.bpj.2013.01.006.

[20] J. P. Butler, I. M. Tolić-Nørrelykke, B. Fabry, and J. J. Fredberg, “Traction fields, moments, and strain energy that cells exert on their surroundings,” Am. J. Physiol. Cell Physiol., vol. 282, no. 3, pp. C595–605, Mar. 2002, doi: 10.1152/ajpcell.00270.2001.

[21] A. Boudaoud et al., “FibrilTool, an ImageJ plug-in to quantify fibrillar structures in raw microscopy images,” Nat. Protoc., vol. 9, no. 2, pp. 457–463, Feb. 2014, doi: 10.1038/nprot.2014.024.

[22] C. Yang et al., “Spatially patterned matrix elasticity directs stem cell fate,” Proc. Natl. Acad. Sci. U. S. A., vol. 113, no. 31, pp. E4439–4445, Aug. 2016, doi: 10.1073/pnas.1609731113.

[23] S. K. Kureel et al., “Soft substrate maintains proliferative and adipogenic differentiation potential of human mesenchymal stem cells on long-term expansion by delaying senescence,” Biol. Open, vol. 8, no. 4, p. bio039453, Apr. 2019, doi: 10.1242/bio.039453.

[24] P. S. Mathieu and E. G. Loboa, “Cytoskeletal and focal adhesion influences on mesenchymal stem cell shape, mechanical properties, and differentiation down osteogenic, adipogenic, and chondrogenic pathways,” Tissue Eng. Part B Rev., vol. 18, no. 6, pp. 436–444, Dec. 2012, doi: 10.1089/ten.TEB.2012.0014.

[25] J. R. Davie, “Inhibition of Histone Deacetylase Activity by Butyrate,” J. Nutr., vol. 133, no. 7, pp. 2485S–2493S, Jul. 2003, doi: 10.1093/jn/133.7.2485S.

[26] C. M. Kraning-Rush, S. P. Carey, J. P. Califano, B. N. Smith, and C. A. Reinhart-King, “The role of the cytoskeleton in cellular force generation in 2D and 3D environments,” Phys. Biol., vol. 8, no. 1, p. 015009, 2011, doi: 10.1088/1478-3975/8/1/015009.

[27] B. R. Sarangi et al., “Coordination between Intra- and Extracellular Forces Regulates Focal Adhesion Dynamics,” Nano Lett., vol. 17, no. 1, pp. 399–406, Jan. 2017, doi: 10.1021/acs.nanolett.6b04364.

[28] “Frontiers | Sodium Valproate-Induced Chromatin Remodeling.” Accessed: Sep. 23, 2026. [Online]. Available: https://www.frontiersin.org/journals/cell-and-developmental-biology/articles/10.3389/fcell.2021.645518/full

[29] S. S. Hur, J. H. Jeong, M. J. Ban, J. H. Park, J. K. Yoon, and Y. Hwang, “Traction force microscopy for understanding cellular mechanotransduction,” BMB Rep., vol. 53, no. 2, pp. 74–81, Feb. 2020, doi: 10.5483/BMBRep.2020.53.2.308.

[30] S. V. Plotnikov, B. Sabass, U. S. Schwarz, and C. M. Waterman, “Chapter 20 - High-Resolution Traction Force Microscopy,” in Quantitative Imaging in Cell Biology, vol. 123, J. C. Waters and T. Wittman, Eds., in Methods in Cell Biology, vol. 123., Academic Press, 2014, pp. 367–394. doi: 10.1016/B978-0-12-420138-5.00020-3.

[31] C. N. Holenstein, U. Silvan, and J. G. Snedeker, “High-resolution traction force microscopy on small focal adhesions - improved accuracy through optimal marker distribution and optical flow tracking,” Sci. Rep., vol. 7, no. 1, p. 41633, Feb. 2017, doi: 10.1038/srep41633.

[32] H. Colin-York et al., “Super-Resolved Traction Force Microscopy (STFM),” Nano Lett., vol. 16, no. 4, pp. 2633–2638, Apr. 2016, doi: 10.1021/acs.nanolett.6b00273.

[33] T. Kampourakis, Y.-B. Sun, and M. Irving, “Myosin light chain phosphorylation enhances contraction of heart muscle via structural changes in both thick and thin filaments,” Proc. Natl. Acad. Sci., vol. 113, no. 21, pp. E3039–E3047, May 2016, doi: 10.1073/pnas.1602776113.

[34] N. Hino et al., “ERK-Mediated Mechanochemical Waves Direct Collective Cell Polarization,” Dev. Cell, vol. 53, no. 6, pp. 646–660.e8, Jun. 2020, doi: 10.1016/j.devcel.2020.05.011.

[35] R. Olivares-Navarrete et al., “Substrate Stiffness Controls Osteoblastic and Chondrocytic Differentiation of Mesenchymal Stem Cells without Exogenous Stimuli,” PLOS ONE, vol. 12, no. 1, p. e0170312, Jan. 2017, doi: 10.1371/journal.pone.0170312.

[36] L. R. Smith, S. Cho, and D. E. Discher, “Stem Cell Differentiation is Regulated by Extracellular Matrix Mechanics,” Physiology, vol. 33, no. 1, pp. 16–25, Jan. 2018, doi: 10.1152/physiol.00026.2017.

[37] A. J. Engler, S. Sen, H. L. Sweeney, and D. E. Discher, “Matrix Elasticity Directs Stem Cell Lineage Specification,” Cell, vol. 126, no. 4, pp. 677–689, Aug. 2006, doi: 10.1016/j.cell.2006.06.044.

[38] J. Oliver-De La Cruz et al., “Substrate mechanics controls adipogenesis through YAP phosphorylation by dictating cell spreading,” Biomaterials, vol. 205, pp. 64–80, Jun. 2019, doi: 10.1016/j.biomaterials.2019.03.009.

[39] E. D. Rosen et al., “PPARγ Is Required for the Differentiation of Adipose Tissue In Vivo and In Vitro,” Mol. Cell, vol. 4, no. 4, pp. 611–617, Oct. 1999, doi: 10.1016/S1097-2765(00)80211-7.

[40] M. V. C. Flores, E. Y. N. Lam, K. E. Crosier, and P. S. Crosier, “Osteogenic transcription factor Runx2 is a maternal determinant of dorsoventral patterning in zebrafish,” Nat. Cell Biol., vol. 10, no. 3, pp. 346–352, Mar. 2008, doi: 10.1038/ncb1697.

[41] W. C. Lee et al., “Multivariate biophysical markers predictive of mesenchymal stromal cell multipotency,” Proc. Natl. Acad. Sci., vol. 111, no. 42, pp. E4409–E4418, Oct. 2014, doi: 10.1073/pnas.1402306111.

[42] S. Dupont, “Role of YAP/TAZ in cell-matrix adhesion-mediated signalling and mechanotransduction,” Exp. Cell Res., vol. 343, no. 1, pp. 42–53, Apr. 2016, doi: 10.1016/j.yexcr.2015.10.034.

[43] J.-H. Hong et al., “TAZ, a transcriptional modulator of mesenchymal stem cell differentiation,” Science, vol. 309, no. 5737, pp. 1074–1078, Aug. 2005, doi: 10.1126/science.1110955.

[44] J.-H. Hwang et al., “Extracellular Matrix Stiffness Regulates Osteogenic Differentiation through MAPK Activation,” PLOS ONE, vol. 10, no. 8, p. e0135519, Aug. 2015, doi: 10.1371/journal.pone.0135519.

[45] P. E. Farahani, S. B. Lemke, E. Dine, G. Uribe, J. E. Toettcher, and C. M. Nelson, “Substratum stiffness regulates Erk signaling dynamics through receptor-level control,” Cell Rep., vol. 37, no. 13, p. 110181, Dec. 2021, doi: 10.1016/j.celrep.2021.110181.

[46] “Mesenchymal Stem Cells for Regenerative Medicine.” Accessed: Sep. 11, 2026. [Online]. Available: https://www.mdpi.com/2073-4409/8/8/886

[47] M. F. Pittenger, D. E. Discher, B. M. Péault, D. G. Phinney, J. M. Hare, and A. I. Caplan, “Mesenchymal stem cell perspective: cell biology to clinical progress,” Npj Regen. Med., vol. 4, no. 1, p. 22, Dec. 2019, doi: 10.1038/s41536-019-0083-6.

[48] M. Romoli et al., “Valproic Acid and Epilepsy: From Molecular Mechanisms to Clinical Evidences,” Curr. Neuropharmacol., vol. 17, no. 10, pp. 926–946, 2019, doi: 10.2174/1570159X17666181227165722.

[49] “Direct Conversion of Human Fibroblasts into Osteoblasts Triggered by Histone Deacetylase Inhibitor Valproic Acid.” Accessed: Sep. 11, 2026. [Online]. Available: https://www.mdpi.com/2076-3417/10/20/7372

