## Supplementary File for "Chromatin Acetylation Overrides Substrate Stiffness-Dependent Mechanoresponse and Lineage Specification in Human Mesenchymal Stem Cells"

**SUPPLEMENTARY INFORMATION**

Table S1

| Acrylamide from 40% stock solution (mL) | Bis-acrylamide from 2% Stock solution (mL) | Water (mL) | E ± St. Dev. (kPa) |
|---|---|---|---|
| 1 | 1.125 | 7.875 | 3.13 ± 0.42 |
| 2.5 | 1.5 | 6 | 34.88 |

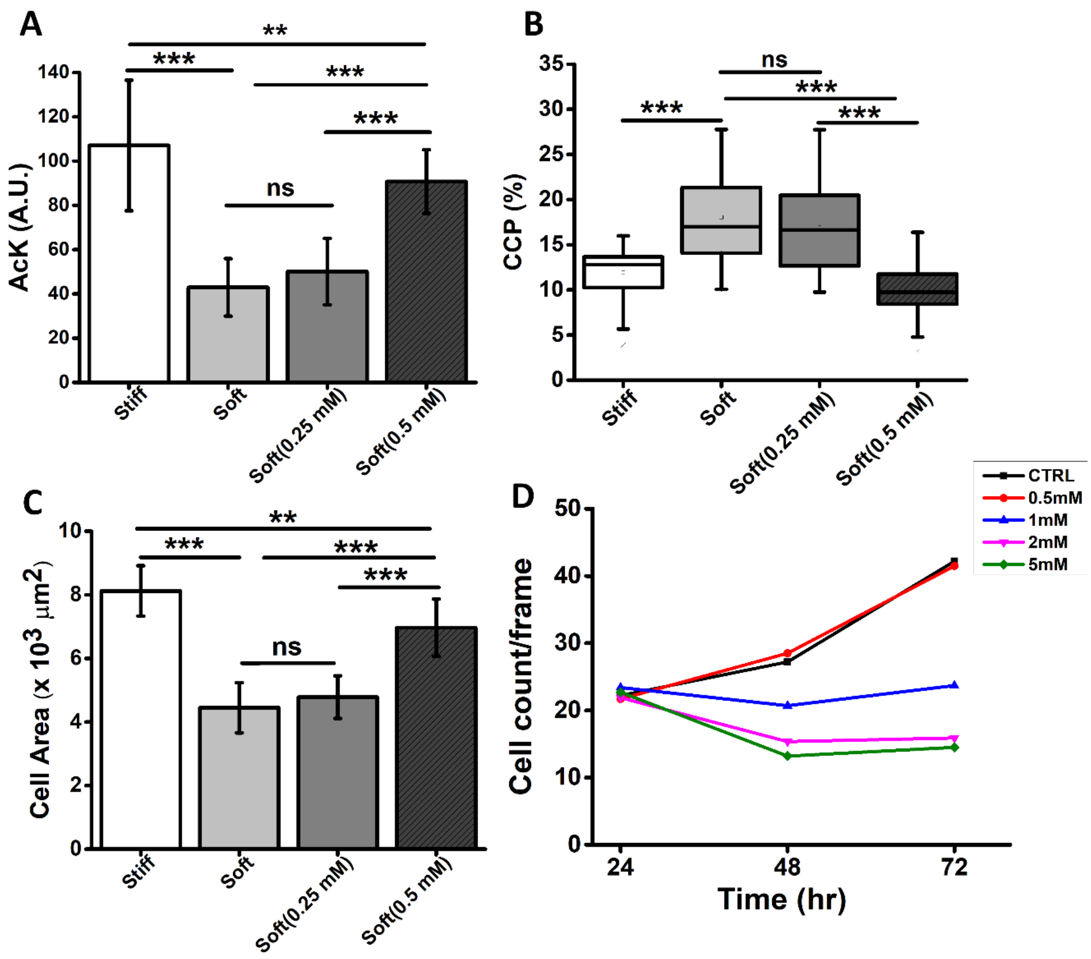


**Fig. S1. Effect of substrate stiffness and varying concentration of VA (HDACi) on spread area, acetylation and CCP.** Quantification showing change in (A) Acetylation (B) chromatin condensation parameter (C) cell spread area. Graph shows significant change in cell area, AcK and CCP at 0.5 mM but no change at lower concentration (0.25 mM). Graph (D) shows at 0.5 mM of HDACi there is no effect on cell growth as compared to control. *p<0.05, **p<0.01, ***p<0.001, N=3,n=50.

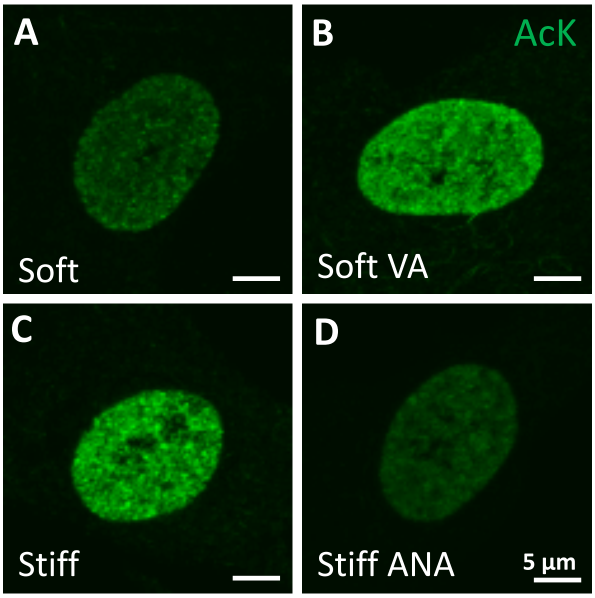


**Fig. S2. Influence of substrate stiffness and HDACi/HATi on chromatin condensation**. (A-B) Representative zoomed-in immunostained images showing intensity of acetylation of lysin residue in histones shown in green for cells cultured on soft (A), soft + VA (B), stiff (C) and stiff +ANA (D); scale bar: 5 μm.

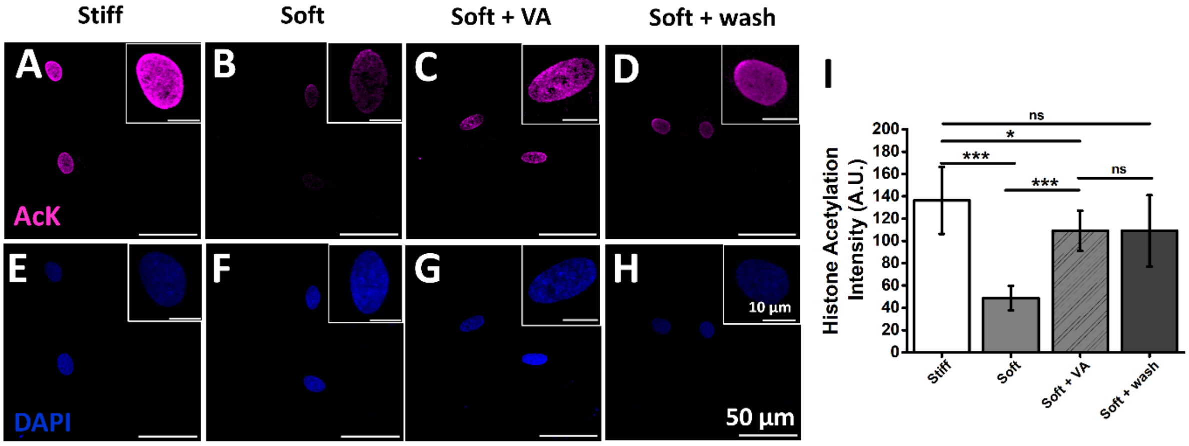


**Fig. S3. Effect of HDACi washing on histone acetylation:** (A-H) Representative immunostained images showing average intensity of acetylation of lysine residue in histones shown in magenta and respective nucleus (DAPI-blue) for cells cultured on stiff (34 kPa, A and E), soft (3 kPa, B and F), soft + VA (C and G) and soft + wash (D and H). Quantification of the same is shown in Fig. I. Our result shows that even after 24 h of washing of VA, the chromatin remains in the open state. *p<0.05, **p<0.01, ***p<0.001, N=3, n>30. scale bar: 50 μm (H) & inset: 10 μm (H).

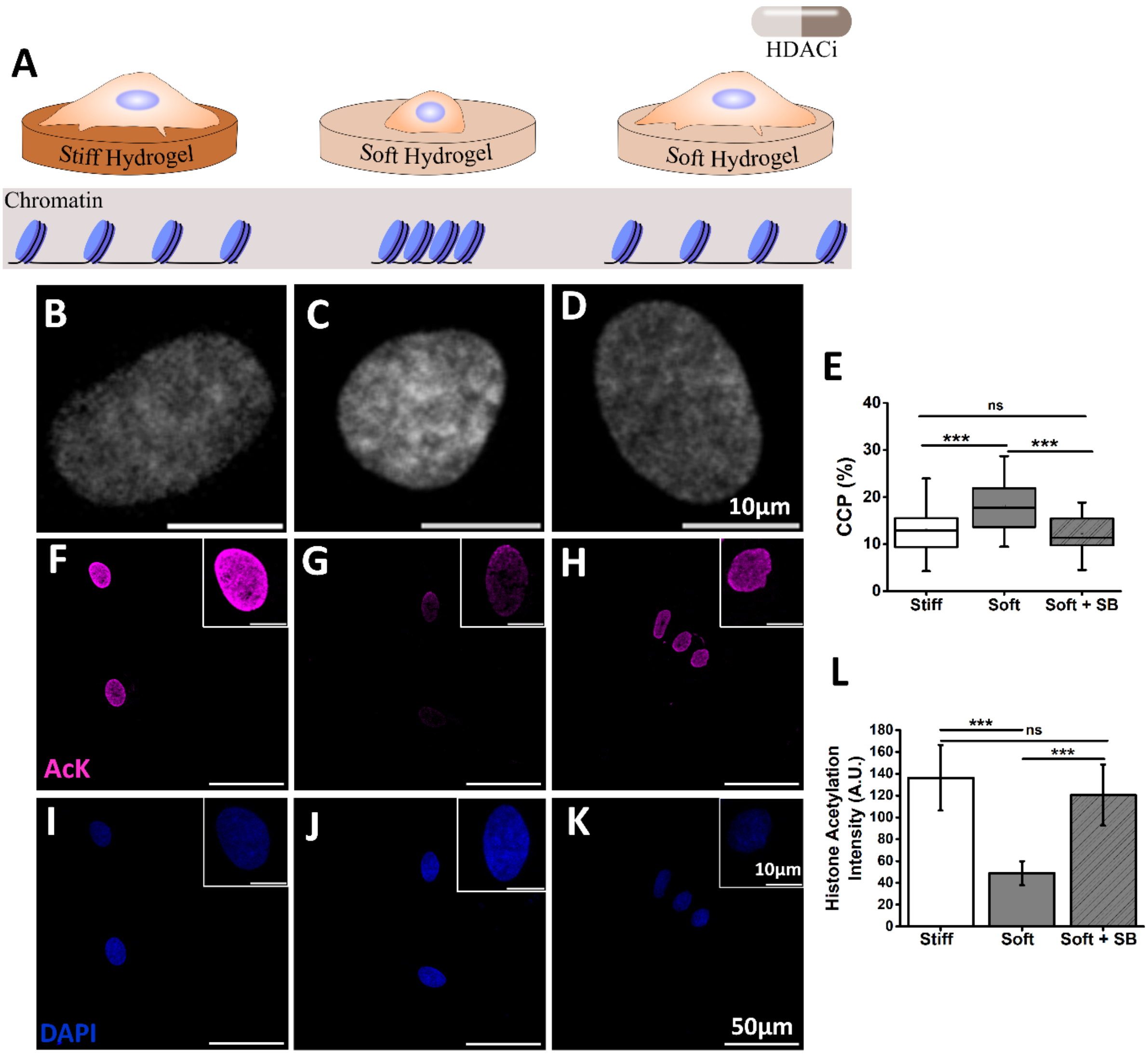


**Fig. S4. Influence of substrate stiffness and sodium butyrate (HDACi) on chromatin condensation**. (A) A schematic showing changes in chromatin condensation due to substrate stiffness and HDACi (Sodium Butyrate, SB). (B-D) Representative images of DAPI-stained nuclei from stiff (34 kPa), soft (3 kPa) and soft + SB. (E) Quantification of CCP defined as % of highly DAPI stained regions to the total nuclear area shown for nuclei from stiff, soft and soft + SB. The result demonstrates that while soft substrate increases chromatin condensation, the same can be brought down to the level of stiff substrates by treating the cells with SB. (F-K) Representative immunostained images showing average intensity of acetylation of lysine residue in histones shown in magenta and respective nucleus (DAPI-blue) for cells cultured on stiff (34 kPa, F and I), soft (3 kPa, G and J) and soft + SB (H and K). Quantification of the same is shown in Fig. L. *p<0.05, **p<0.01, ***p<0.001, N=3, n>30. scale bar: 10 µm (D), 50 µm (K) & inset: 10 µm (K).

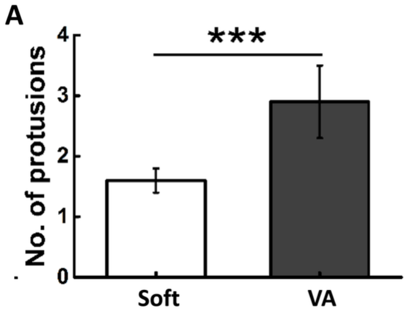


**Fig. S5. Effect of HDACi on focal adhesion area and actin anisotropy.** Graph (A) show quantification of no. of protrusions in each cell when cultured on soft gel with and without VA. *p<0.05, **p<0.01, ***p<0.001.

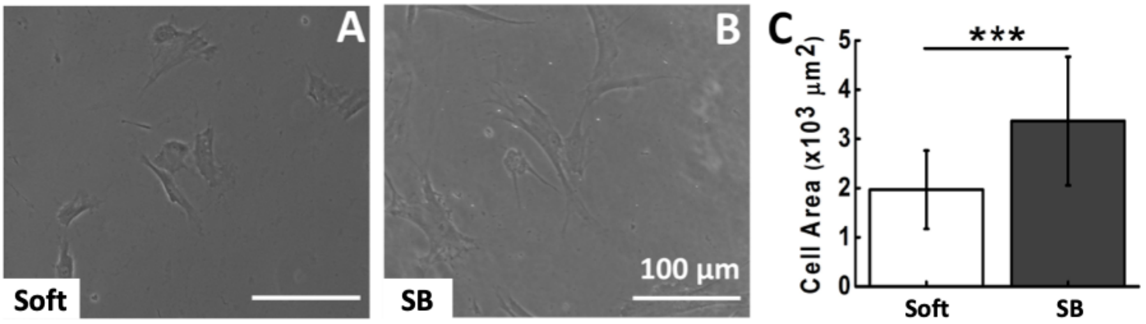


**Fig. S6. HDACi increases cell spread area on soft substrate.** Representative images of hMSCs cultured on 3 kPa gel (A) without SB and (B) with SB. Graph (C) show quantitative analysis of change in projected cell area on 3 kPa gels with and without SB. **p*<0.05, **p<0.01, ***p<0.001, N=3, n > 50 cells, scale bar: 100 µm.

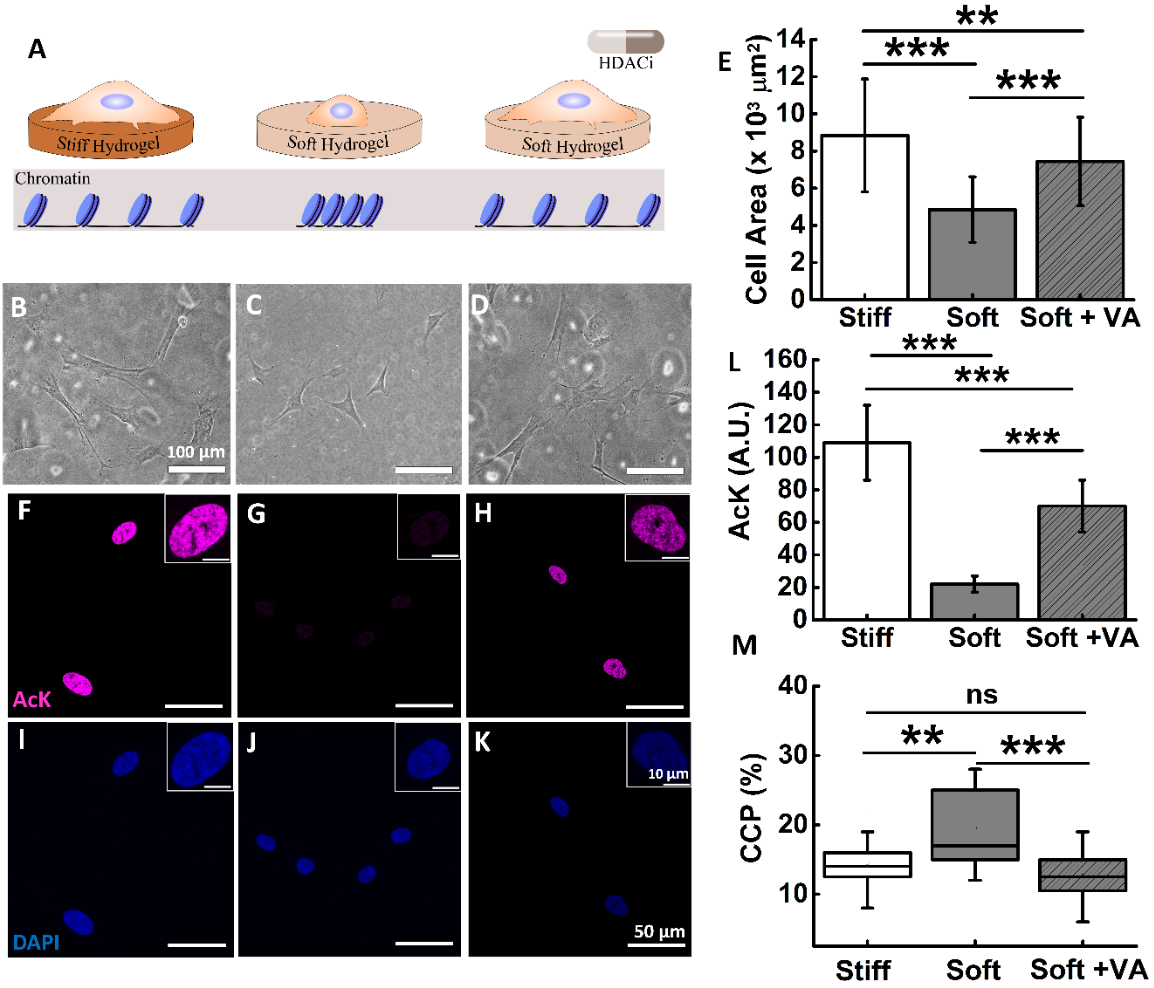


**Fig. S7. Influence of fibronectin coating on hMSCs morphology and chromatin condensation in presence of HDACi on stiff and soft substrates.** (A) schematic showing changes in cellular morphology and chromatin condensation due to substrate stiffness and HDACi (VA). (B-D). Representative phase contrast images of hMSCs cultured on stiff (34 kPa), soft (3 kPa) and soft + VA. (F-H) Representative images of acetylation (magenta) and (I-K) respective nucleus (DAPI-blue) for cells cultured on stiff (34 kPa), soft (3 kPa) and soft + VA. Graph (E) shows quantification of cell spread area of hMSCs cultured on stiff (34 kPa), soft (3 kPa) and soft + VA. Graph ( L & M) shows quantification of average acetylation intensity and CCP of cells cultured on stiff (34 kPa), soft (3 kPa) and soft + VA. *p<0.05, **p<0.01, ***p<0.001, N=3, n>50, scale bar: 100 µm (B), 50 µm (K), inset : 10µm (K).

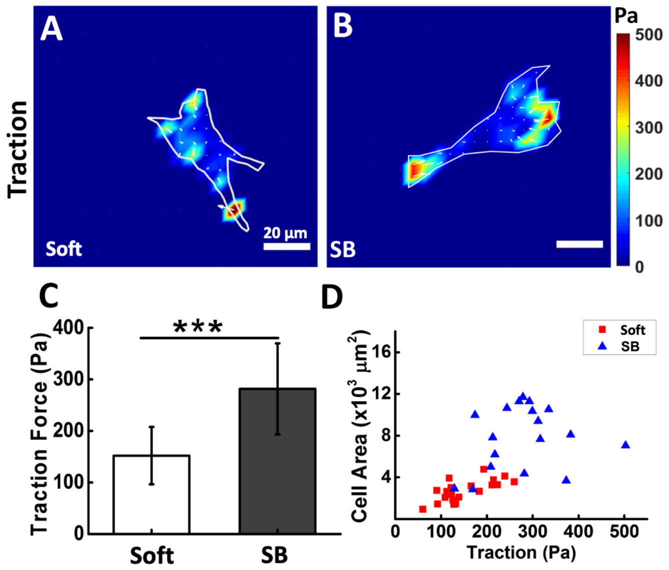


**Fig. S8. HDACi, sodium butyrate increases hMSCs traction on soft substrates.** Heat map of soft substrate-cultured hMSCs traction stress (A) without SB, (B) with SB. Colours corresponds to magnitude of stresses as indicated in the colour bar. Graph (C) shows quantitative analysis of traction force on 3 kPa gel with and without SB (Ctrl). Scatter plot (D) shows positive correlation between cell spread area and cellular traction. *$p$<0.05, **p<0.01, ***p<0.001, N=2, n>15 cells, scale bar: 20 µm.

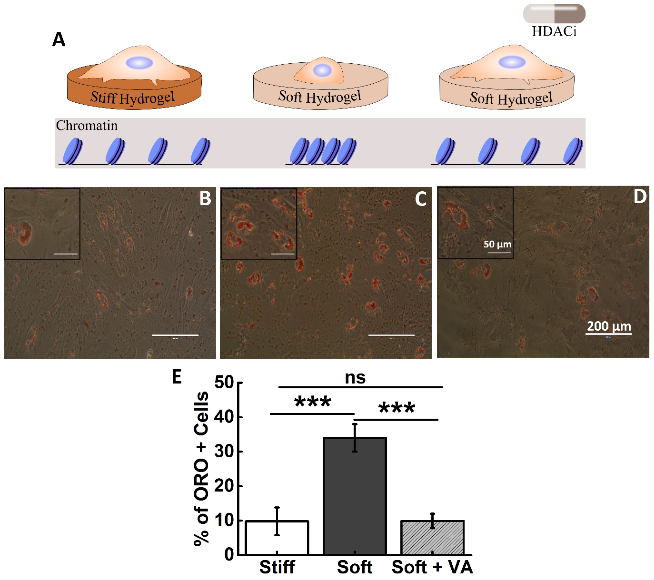


**Fig. S9. HDACi overrides the adipogenic differentiation potential of hMSCs cultured on soft substrates (3 kPa).** (A) schematic showing changes in chromatin condensation due to substrate stiffness and HDACi**.** Representative images of Oil Red O staining for hMSCs cultured on stiff (34 kPa), soft (3 kPa) and soft + VA. The insets are the magnified local image to show Oil Red O formation. Graph (E) shows quantitative analysis of change in percentage of ORO positive cells for hMSCs cultured on stiff (34 kPa), soft (3 kPa) and soft + VA. *p<0.05, **p<0.01, ***p<0.001, N=3, n=300, scale bar: 200 µm, inset : 50 µm.

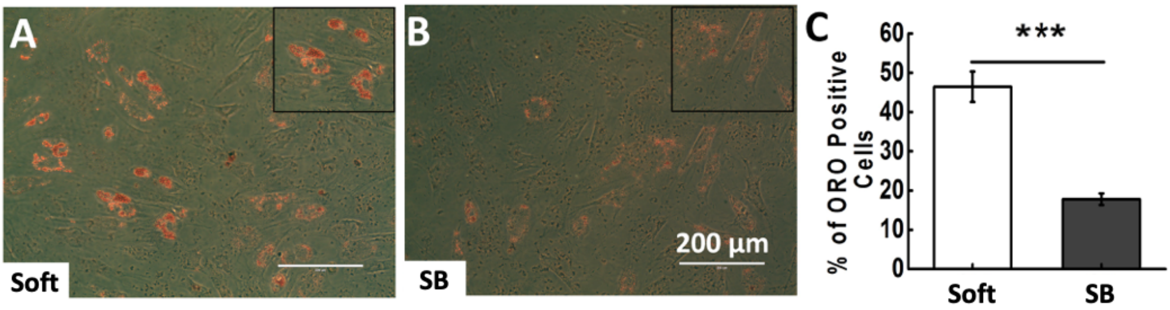


**Fig. S10. HDACi inhibit adipogenesis of hMSCs cultured on soft substrates.** Representative images of Oil Red O staining on 3 kPa gel without SB (A) and with SB (B). The insets are the magnified local image to show oil droplets. Graph (C) shows quantitative analysis of change in percentage of ORO positive cells with and without SB. **p*<0.05, **p<0.01, ***p<0.001, N=3, n=300, scale bar: 200 µm.

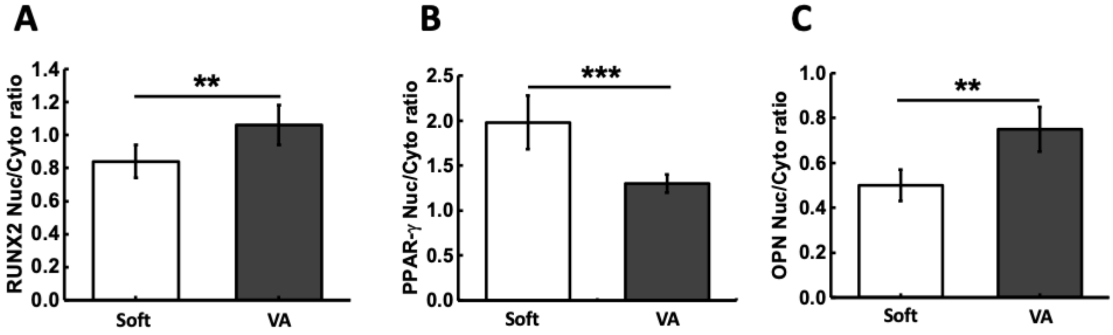


**Fig. S11. Effect of HDACi on nuclear/cytoplasmic ratio of differentiation markers.** Graph (A), (B) and (C) show quantification of RUNX2, PPAR-γ and osteopontin nuclear to cytoplasmic ratio respectively, when the cells are cultured on soft gel with and without VA. *p<0.05, **p<0.01, ***p<0.001, N=3, n>25.

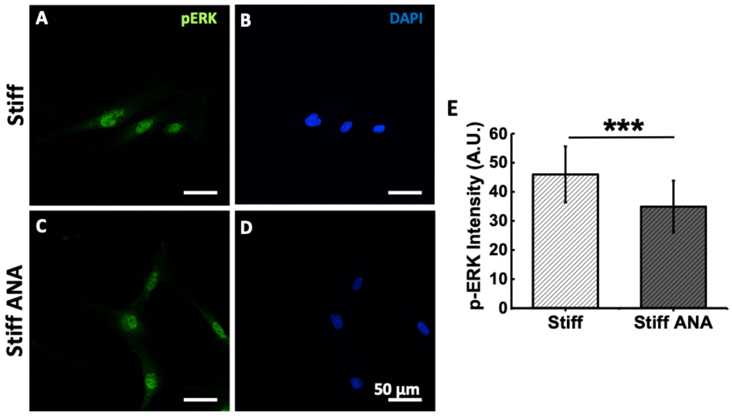


**Fig. S12. Effect of HATi on pERK expression on stiff substrate.** (A-D) Representative p-ERK (green) and DAPI (blue) stained immunofluorescence images has been shown for hMSCs cultured on 34 kPa substrate without and with ANA respectively. (G) The graph of quantitative measurement of p-ERK intensity in hMSCs has been shown. *p<0.05, **p<0.01, ***p<0.001, N=3, n>25.

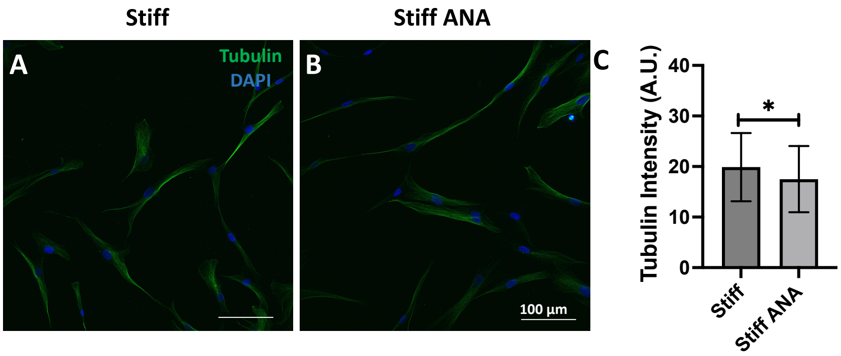


**Fig. S13. Effect of HATi on Tubulin expression on stiff substrate.** (A and B) Representative immunofluorescence images of Tubulin (green) and DAPI (blue), along with the corresponding merged panels, for hMSCs cultured on 34 kPa substrates without and with ANA, respectively. (C) The quantitative measurement of Tubulin intensity in hMSCs has been shown. *p<0.05, n>50.

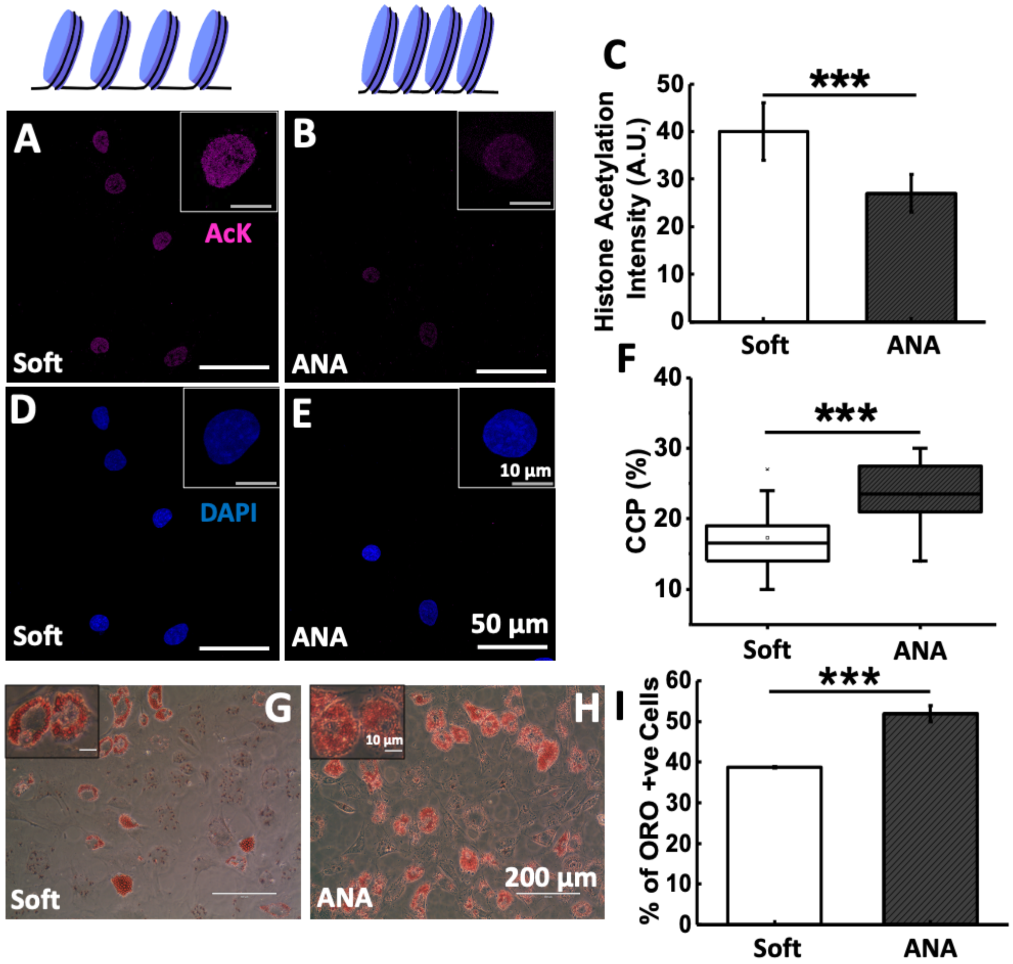


**Fig. S14. Chromatin condensation promotes adipogenesis of hMSCs cultured on soft substrate:** (A-B) Representative images of acetylation (magenta) and (D-E) respective nucleus (DAPI-blue) from Soft (3 kPa) (A and D) and Soft + ANA (B and E). (C) shows corresponding quantification of Histone acetylation average intensity. (F) Shows quantification of % CCP for the nuclei from Soft (3 kPa) and Soft + ANA. Oil Red O staining after ten days of culturing hMSCs in adipogenic induction media (AIM) on soft gel (3 kPa) without (G) and (H) with ANA. (I) Quantification of the percentage of lipid oil droplets in the presence of AIM and AIM with ANA shows that treatment with ANA further increases adipogenesis on soft gel. *$p<0.05$, **$p<0.01$, ***$p<0.001$, N=3, n>30. Scale bar:50 µm (E), inset:10 µm (E), 200 µm (H), inset:10 µm (H).
